# Fast bacterial growth reduces antibiotic accumulation and efficacy

**DOI:** 10.1101/2021.10.18.464851

**Authors:** Urszula Łapińska, Margaritis Voliotis, Ka Kiu Lee, Adrian Campey, M. Rhia L. Stone, Wanida Phetsang, Bing Zhang, Krasimira Tsaneva-Atanasova, Mark A. T. Blaskovich, Stefano Pagliara

**Author notes:** Stefano Pagliara & Urszula Łapińska, **Email:**.

## Abstract

Phenotypic variations between individual microbial cells play a key role in the resistance of microbial pathogens to pharmacotherapies. Nevertheless, little is known about cell individuality in antibiotic accumulation. Here we hypothesize that phenotypic diversification can be driven by fundamental cell-to-cell differences in drug transport rates. To test this hypothesis, we employed microfluidics-based single-cell microscopy, libraries of fluorescent antibiotic probes and mathematical modelling. This approach allowed us to rapidly identify phenotypic variants that avoid antibiotic accumulation within populations of *Escherichia coli, Pseudomonas aeruginosa, Burkholderia cenocepacia* and *Staphylococcus aureus*. Crucially, we found that fast growing phenotypic variants avoid macrolide accumulation and survive treatment without genetic mutations. These findings are in contrast with the current consensus that cellular dormancy and slow metabolism underlie bacterial survival to antibiotics. Our results also show that fast growing variants display significantly higher expression of ribosomal promoters before drug treatment compared to slow growing variants. Drug-free active ribosomes facilitate essential cellular processes in these fast growing variants, including efflux that can reduce macrolide accumulation. Using this new knowledge, we phenotypically engineered bacterial populations by eradicating variants that displayed low antibiotic accumulation through the chemical manipulation of their outer membrane inspiring new avenues to overcome current antibiotic treatment failures.

## Introduction

Phenotypic heterogeneity between genetically identical cells has been observed across all three domains of life(1,2). This heterogeneity is characterized by individual cells that display differing phenotypic traits(3,4) and permit genotypes to persist in fluctuating environments(2). Phenotypic heterogeneity in the bacterial response to antibiotics contributes to antimicrobial resistance(5–11) and the failure to effectively treat bacterial infections(12–14). Therefore, it is imperative to develop new diagnostics capable of rapidly identifying phenotypic variants that survive antibiotic treatment(15) and develop new antibiotic therapies against such phenotypic variants(16).

Here we hypothesize that this phenotypic diversification is driven by fundamental cell-to-cell differences in membrane transport mechanisms and their underpinning regulatory networks. In order for an antibiotic to be effective, it needs to reach its cellular target at a concentration that is inhibitory for microorganism growth(17). In gram-negative bacteria, intracellular antibiotic accumulation(17–19) is a complex biophysical phenomenon involving different physicochemical pathways and a combination of exquisitely regulated active and passive transport processes(17,20). These processes include diffusion through the outer membrane lipid bilayer(17) and porins(21,22); self-promoted uptake through the outer membrane(23); diffusion through the inner membrane lipid bilayer which displays orthogonal selection properties compared to the outer membrane(24,25); active transport via inner membrane transporters(24); efflux out of the cell(26–29); enzymatic modification or degradation(17); and eventually binding to the intracellular target.

Learning the rules that permit antibiotics to accumulate in gram-negative bacteria is vitally important in order to combat phenotypic and genotypic resistance to antibiotics(24,30,31). However, most permeability data are sequestered in proprietary databases(17). Moreover, such experimental datasets have often been generated via cell-free methods that permit the measurement of the diffusion rate of a compound through simplified membrane pathways(32), but care should be taken when projecting these data to the more complex accumulation dynamics in live cells(17). Live or fixed cell methodologies including radiometric, fluorometric or biochemical assays(33–35), mass spectrometry(36–42), Raman spectroscopy(43) and microspectroscopy(44–46) have also been employed to carry out antibiotic accumulation assays. These techniques generally rely on ensemble measurements that average the results obtained from a large population of microorganisms, or are derived from examining only a handful of individual bacteria. Therefore, little is known about the variability in individual drug accumulation across many single cells within a clonal population.

Here, we fill this fundamental gap in our knowledge by harnessing the power of microfluidics-microscopy(47,48) combined with fluorescent antibiotic-derived probes(49–51) as well as unlabelled antibiotics. This approach allows us to examine the interactions between the major classes of antibiotics and hundreds of live individual bacteria in real-time whilst they are being dosed with the drugs. Combined with mathematical modelling these data allow us to rapidly identify phenotypic variants that avoid antibiotic accumulation and are able to sustain growth in the presence of drugs without acquiring genetic mutations. We show that bacteria close to the antibiotic source accumulate faster membrane-targeting antibiotics but more slowly antibiotics with intracellular targets compared to bacteria further away from the antibiotic source. In contrast with the current consensus that slow cell growth leads to reduced antibiotic efficacy, we discover that fast growing phenotypic variants avoid macrolide accumulation due to a higher abundance of both ribosomes (i.e. the drug target) and efflux pumps. We further demonstrate that chemically manipulating the bacterial outer membrane permits us to phenotypically engineer bacterial populations by eradicating variants that display low antibiotic accumulation. Adopting our novel approach in clinical settings to inform the design of improved drug therapies could radically transform our one health approach to antimicrobial resistance.

## Results

### Experimental assessment of single-cell real-time drug accumulation dynamics

We combined our recently developed single-cell microfluidics-microscopy platform(47,48,52) with a library of fluorescent derivatives representing most major classes of antibiotics, including macrolides (roxithromycin)(52), oxazolidinones (linezolid)(53), glycopeptides (vancomycin)(54), fluoroquinolones (ciprofloxacin)(55), antifolates (trimethoprim)(56), and membrane-targeting lipopeptides/peptides (polymyxin B, octapeptin, tachyplesin)(54) (Fig. 1A).

**Figure 1.**
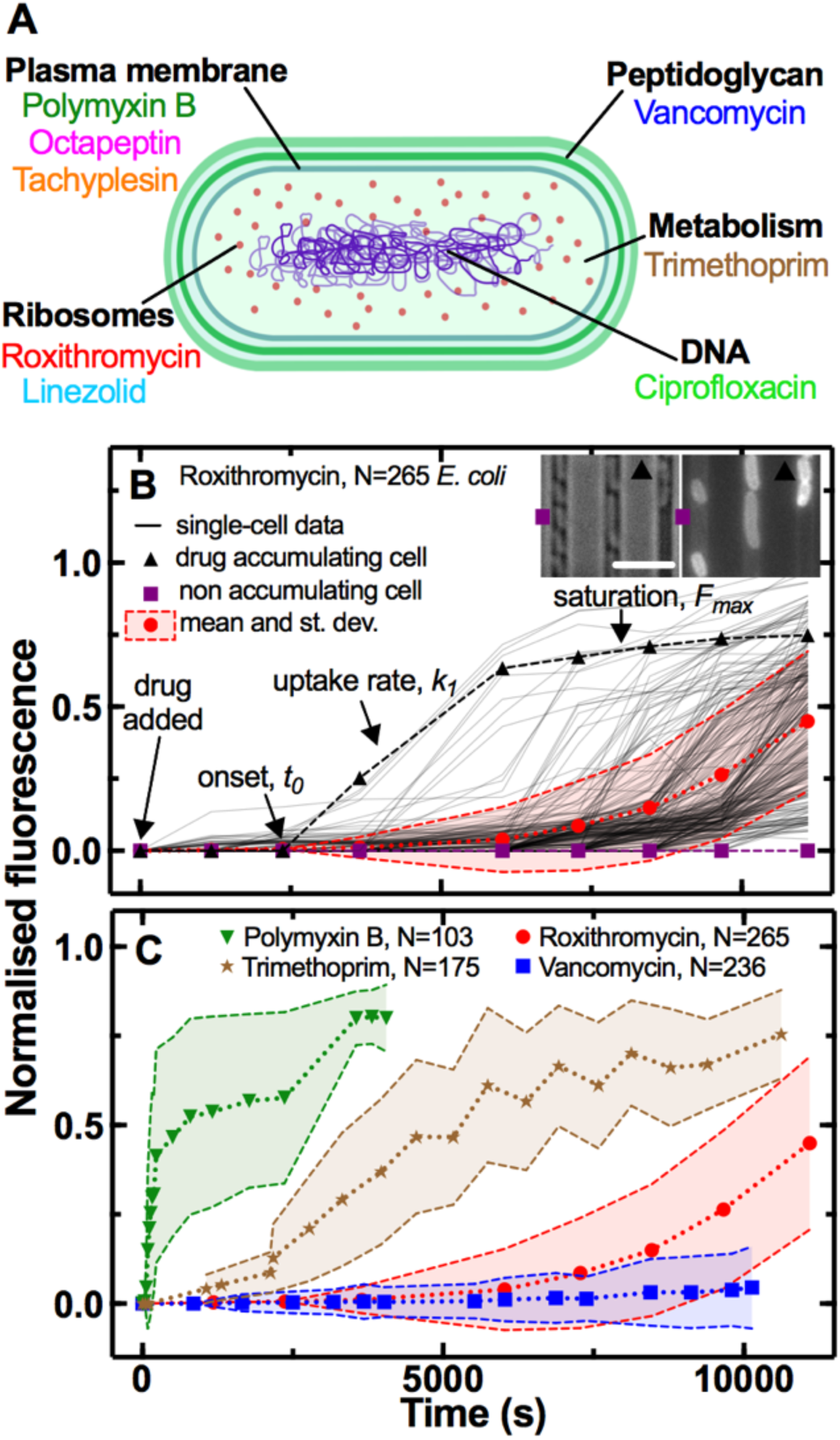
Phenotypic heterogeneity in the accumulation of the major classes of antibiotics. **A**) Illustration depicting the eight antibiotics employed in this study alongside their bacterial targets. **B**) Accumulation of the fluorescent derivative of roxithromycin in 265 individual *E. coli* (continuous lines) after adding the probe at 46 μg mL^-1^ extracellular concentration in M9 minimal medium from t=0 onwards. Fluorescence values were background subtracted and normalised first by cell size and then to the maximum value in the dataset (see Methods). The circles and shaded areas represent the mean and standard deviation of the values from 265 bacteria collated from biological triplicate. The squares represent the fluorescent values of a representative bacterium that does not accumulate the fluorescent derivative of roxithromycin, whereas the triangles represent the fluorescent values of a representative bacterium that accumulates the drug. Insets: representative brightfield and fluorescence images after 7,000 s incubation in the fluorescent derivative of roxithromycin, the symbols indicate the two representative bacteria above. Scale bar: 5 μm. *t*_*0*_, *k*_*1*_ and *F*_*max*_ indicate the time point at which single-cell fluorescence becomes distinguishable from the background, the rate of uptake and the final levels of accumulation at steady-state, respectively. **C**) Population average (symbols) and standard deviation (shaded areas) of the accumulation of the fluorescent derivatives of polymyxin B (triangles), trimethoprim (stars), roxithromycin (circles) and vancomycin (squares) probes added at 46 μg mL^-1^ extracellular concentration in M9 minimal medium from t=0 onwards. Data are obtained by averaging at least one hundred single-cell values (i.e. N=103, 175, 265 and 236, respectively) collated from biological triplicate. Corresponding single-cell data along with data for the fluorescent derivatives of linezolid, tachyplesin, octapeptin and ciprofloxacin probes are reported in Fig. S1.

Each antibiotic was functionalised at a site that minimises any changes in biological activity, adding a substituent that allows for facile coupling with a small fluorophore, nitrobenzoxadiazole (NBD, Table S1) as previously reported(52–56). We confirmed that the majority of fluorescent derivatives retained the antibiotic activity of the parent drug via minimum inhibitory concentration (MIC) assays (Table S1). Next we used each probe in our microfluidics-microscopy platform(47,48,52) to quantify the dynamics of the accumulation of each antibiotic in individual bacteria in real-time (Fig. 1B). Briefly, we loaded an aliquot of a stationary phase clonal bacterial culture in a microfluidic device equipped with small parallel channels, each hosting between one to six bacteria(47,48,52). Then we continuously flowed lysogeny broth (LB) medium into the device for 2 h to stimulate cell growth and reproduction. At this point, we injected one of the antibiotic probes and imaged the real-time intracellular probe accumulation in hundreds of individual live bacteria (Video S1 and S2). Typically, upon onset (*t*_*0*_), the uptake was initially linear (with rate constant *k*_*1*_), before reaching steady-state saturation levels (*F*_*max*_, Fig. 1B) due to probe efflux, compound transformation(17), or target saturation(24), although several bacteria displayed divergent accumulation dynamics (Fig. 1B, S1 and S2).

### Heterogeneity in antibiotic accumulation in gram-negative and gram-positive bacteria

These single-cell measurements revealed hitherto unrecognised phenotypic heterogeneity in intracellular drug accumulation in clonal populations of *E. coli* as evident from the microscopy images in Fig. 1B and Fig. S1. In contrast, standard techniques measure population averages of drug accumulation across thousands or millions of cells(19,33,35–39,42). In our single-cell assay, population averages (circles in Fig. 1B) did not reflect the fact that some phenotypic variants displayed a remarkably delayed onset, slower uptake rate or reduced saturation with respect to other cells (e.g. compare the accumulation trajectories reported by the squares - no accumulation - vs triangles - high accumulation - in Fig. 1B). These phenotypic variants have thus far remained unrecognised in population-based experiments and give rise to large coefficients of variation (CV, the ratio of the standard deviation over the mean) in the accumulation of each of the eight investigated antibiotics (Fig. 1C and S3). In the following we will therefore use CV as a reporter for phenotypic heterogeneity within bacterial populations as previously reported(57).

All bacteria within each experiment were exposed to the same concentration of probe (46 μg mL^-1^) for the same duration and to the same drug milieu, i.e. minimal medium M9 to avoid dilution of probes due to cell growth(17). In accordance with previous studies about phenotypic responses to antibiotics(58,59), we found that bacterial variants displaying delayed or reduced antibiotic accumulation were genuine phenotypic variants, since DNA sequencing of the device outflow did not reveal any genetic mutations compared to untreated bacteria. Furthermore, these variants did not display significant differences in cell size (Fig. S4) and we further normalised each single-cell fluorescence value to the corresponding single-cell size (see Methods)(60).

Due to the presence of these phenotypic variants, not all the bacteria were stained by each antibiotic probe, thus we found drug-dependent dynamics in the fraction of stained bacteria (Fig. S5). The lipopeptide/peptide probes targeting the outer bacterial membrane (polymyxin B, octapeptin and tachyplesin) stained 90% of the investigated bacteria within 1,000 s post-addition to the microfluidic device. At this time, the trimethoprim and ciprofloxacin probes targeting intracellular components had stained only 50% of the bacteria, whereas the number of bacteria stained by roxithromycin and vancomycin probes, with a large molecular weight (1064 and 1650 g mol^-1^, respectively), was close to zero. However, the roxithromycin probe did stain 50% and 90% of the bacteria around 7,500 s and 9,000 s, respectively, post-addition to the device, by which time only 15% of the bacteria had been stained by vancomycin. The lack of vancomycin staining was expected since vancomycin cannot cross the gram-negative double membrane to access its peptidoglycan target(61).

Next, we verified that this hitherto unrecognised heterogeneity in antibiotic accumulation is not a phenotypic feature exclusive to *E. coli*. When we compared and contrasted roxithromycin-NBD accumulation in *E. coli* against uptake in the gram-positive bacterium *S. aureus*, we found that although the latter displayed more rapid accumulation dynamics (Fig. S6A and S6B, respectively), also *S. aureus* displayed phenotypic variants with delayed or reduced accumulation. In fact, roxithromycin-NBD reached saturation levels 3,000 s post-addition in some *S. aureus* cells, whereas other bacteria accumulated the drug at very low levels and only by 5,000 s post-addition (with a CV in range 53-372% and 29-73% for *E. coli* and *S. aureus*, respectively). In contrast, the gram-positive targeting vancomycin-NBD readily and homogeneously accumulated in *S. aureus* within 2,500 s post-addition (CV in range 12-14%, Fig. S7B), but did not accumulate in *E. coli* (within this same timeframe, Fig. S7A). Finally, we found phenotypic variants with delayed or reduced accumulation of ciprofloxacin-NBD in three clinically-relevant gram-negative bacteria: *E. coli, Pseudomonas aeruginosa* and *Burkholderia cenocepacia* (CV in range 12-329%, 24-534% and 31-90%, Fig. S8A, S8B and S8C, respectively). Furthermore, ciprofloxacin-NBD accumulated more slowly and to a lower extent in *P. aeruginosa* compared to *E. coli* and *B. cenocepacia* (Fig. S8) in accordance with previous measurements at the whole population level(35) and possibly due to the high porin impermeability in *P. aeruginosa*(62).

Finally, in order to verify that neither the drug milieu nor the concentration nor the labelling underpin the observed heterogeneity in antibiotic accumulation, we run separate controls using *E. coli* and both sub-inhibitory and inhibitory concentrations of roxithromycin-NBD dissolved either in M9 or LB (Fig. S9), as well as unlabelled ciprofloxacin, ciprofloxacin-NBD, roxithromycin-NBD and roxithromycin-DMACA (dimethylaminocoumarin-4-acetate, Fig. S10). In all cases we identified phenotypic variants with delayed or reduced antibiotic accumulation, leading to large CVs as shown in Fig. S9 and Fig. S10. We can also exclude possible effects of variations in magnesium availability(23,63) on the measured heterogeneity in antibiotic accumulation since all bacteria were exposed to the same medium within the microfluidic device.

### Single-cell coupling between kinetic accumulation parameters

Prompted by these novel findings, we moved on to an in-depth examination of antibiotic accumulation dynamics and the underlying cellular and molecular mechanisms. Firstly, we developed and implemented a mathematical model to rationalise these markedly heterogeneous single-cell accumulation dynamics, including phenotypic variants with delayed or reduced antibiotic accumulation (see Methods). Briefly, this model describes drug accumulation based on two coupled ordinary differential equations. The first equation describes drug accumulation in terms of uptake, which proceeds at a time-varying rate, and drug loss (due to efflux or degradation(17)), which we assume to be a first order reaction with rate constant *d*_*c*_. The second equation describes how the uptake rate changes over time. Here we assume a state of uptake (parameter *k*_*1*_, which switches on with a time delay; parameter *t*_*0*_); a linear decay term (parameter *d*_*r*_); as well as an adaptive inhibitory effect (parameter *k*_*2*_) of the intracellular drug concentration on the uptake rate (allowing us to capture the dip we observe in some single-cell trajectories in Fig. S2). We used this model to fit our single-cell *E. coli* data on the accumulation of all the above investigated drugs apart from vancomycin. This allowed us to compare and contrast the accumulation kinetic parameters above for the different antibiotics, since we used the same probe concentration for each drug (46 μg mL^-1^) and all drugs were tested against the same clonal *E. coli* population. For vancomycin we found poor fitting for the majority of cells (195 out of 241 cells), as the fluorescent signal remained indistinguishable from the background, due to low cellular uptake (Fig. S1H).

Membrane targeting antibiotic probes displayed on average faster accumulation onset (*t*_*0*_ = 306, 364 and 571 s^-2^ for tachyplesin, polymyxin B and octapeptin, respectively) compared to antibiotics with an intracellular target (*t*_*0*_ = 437, 2,525, 3,608 and 6,614 s^-2^ for linezolid, trimethoprim, ciprofloxacin and roxithromycin, respectively, Fig. S11). Remarkably, we found notable cell-to-cell differences in *t*_*0*_ across all investigated drugs with a maximum CV of 209% for polymyxin B, and a minimum CV of 25% for roxithromycin (Fig. S11), further confirming the presence of phenotypic variants with delayed antibiotic accumulation.

Membrane targeting antibiotic probes also displayed, on average, steeper rates of uptake (*k*_*1*_ = 260, 229 and 93 a.u. s^-2^ for tachyplesin, polymyxin B and octapeptin, respectively) compared to antibiotics with an intracellular target (*k*_*1*_ = 4.4, 1.6, 0.9 and 0.3 a.u. s^-2^ for roxithromycin, linezolid, ciprofloxacin and trimethoprim, respectively, Fig. S11). Also, *k*_*1*_ was heterogeneous across all drugs investigated with a maximum CV of 124% for roxithromycin and a minimum CV of 37% for trimethoprim (Fig. S11), further confirming the presence of phenotypic variants with slow antibiotic uptake.

Membrane targeting antibiotic probes also displayed, on average, higher steady-state saturation levels (*F*_*max*_ = 2,597, 2,357 and 2,264 a.u. for tachyplesin, octapeptin and polymyxin B, respectively) compared to antibiotics with an intracellular target (*F*_*max*_ = 1,034, 512, 253 and 180 a.u. for roxithromycin, linezolid, trimethoprim and ciprofloxacin, respectively, Fig. S11). *F*_*max*_ was also heterogeneous with a maximum CV of 55% for roxithromycin and a minimum CV of 9% for octapeptin (Fig. S11) further confirming the presence of phenotypic variants with reduced antibiotic accumulation. For brevity, the second order kinetic parameters *k*_*2*_, *d*_*r*_, and *d*_*c*_ are reported and discussed only in Fig. S12.

The finding that accumulation of membrane targeting probes happens earlier, faster and to a greater extent than probes with an intracellular target can be easily rationalised considering that the latter probes need to cross the gram-negative double membrane. This represents a very good validation of our combined experimental and theoretical approach. However, the large heterogeneity in the kinetic parameters describing the accumulation of all probes, due to phenotypic variants with delayed or reduced accumulation, was instead unexpected. Additionally, the finding that roxithromycin simultaneously displayed the most delayed accumulation onset but also the steepest rate of uptake and highest steady-state saturation levels, across antibiotic probes with intracellular targets, was also unexpected. These data corroborate the hypothesis that multiple mechanisms must be involved in intracellular antibiotic accumulation at the level of the individual cell(17), a point which we expand on below.

Next, we used the inferred accumulation kinetic parameters to test the hypothesis that phenotypic variants within a clonal population specialise to reduce antibiotic accumulation. When we pooled together the single-cell values for all the antibiotics tested against *E. coli*, we found a strong negative correlation between *t*_*0*_ and *k*_*1*_ and *t*_*0*_ and *F*_*max*_, but a strong positive correlation between *k*_*1*_ and *F*_*max*_ (Fig. 2A-C, Pearson coefficients r = −0.40, −0.27 and 0.65, respectively, p<0.0001; the relationship between *k*_*1*_ and *F*_*max*_ is partially imposed by the definition of *F*_*max*_ in the model, whereas the ones between *t*_*0*_ and *k*_*1*_ or *F*_*max*_ are not).

**Figure 2.**
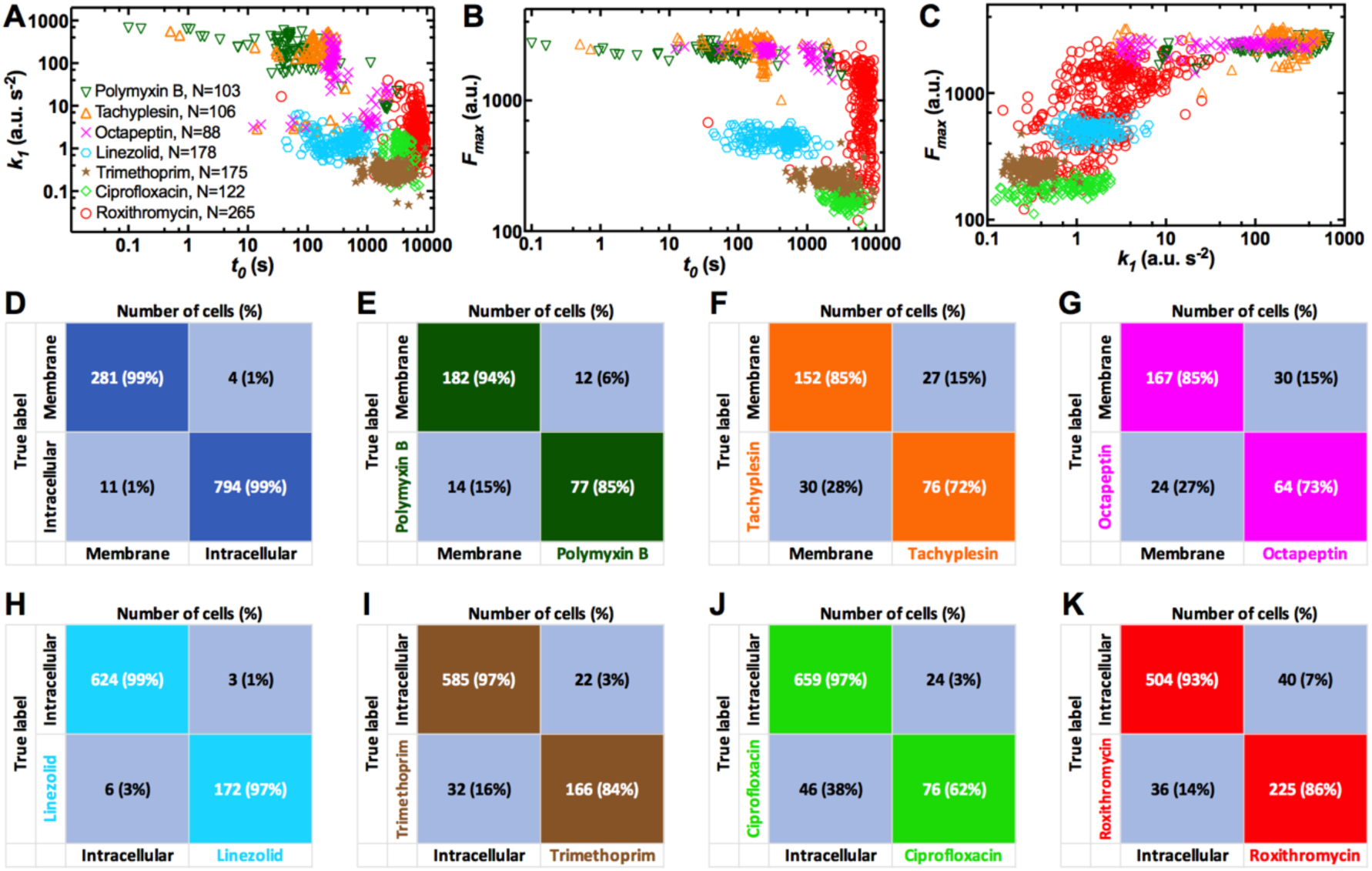
Single-cell coupling between key kinetic accumulation parameters. Correlation between **A**) *t*_*0*_ and *k*_*1*_, **B**) *t*_*0*_ and *F*_*max*_, **C**) *k*_*1*_ and *F*_*max*_ describing the accumulation of the fluorescent derivatives of polymyxin B (downward triangles), tachyplesin (upward triangles), octapeptin (crosses), linezolid (hexagons), trimethoprim (stars), ciprofloxacin (diamonds) or roxithromycin (circles) in N = 103, 106, 88, 178, 175, 122, 265 individual *E. coli*, respectively. Each data point represents the values of two kinetic parameters inferred for an individual bacterium from the data in Fig. S1 using our mathematical model. Statistical classification of the accumulation of **D)** membrane- (i.e. polymyxin B, tachyplesin and octapeptin) vs intracellular-targeting antibiotics (i.e. linezolid, trimethoprim, ciprofloxacin, roxithromycin), **E)** polymyxin B, **F)** tachyplesin or **G)** octapeptin vs the remaining membrane-targeting antibiotics, **H)** linezolid, **I)** trimethoprim, **J)** ciprofloxacin or **K)** roxithromycin vs remaining antibiotics with an intracellular target. These confusion tables are predictions generated using only the two kinetic parameters that can be rapidly measured experimentally, namely *t*_*0*_ and *k*_*1*_. Similar statistical classifications were obtained when using the full set of kinetic parameters, i.e. *k*_*2*_, *d*_*r*_, and *d*_*c*_ in addition to *t*_*0*_ and *k*_*1*_.

These strong correlations show that the bacteria which start accumulating drugs later also display slow uptake and low saturation levels. This statistical analysis also reveals that i) it is possible to rapidly identify phenotypic variants displaying reduced antibiotic accumulation by inferring the whole set of kinetic parameters from a smaller subset (e.g. by inferring *F*_*max*_ from *t*_*0*_ and *k*_*1*_, the latter two can be measured significantly faster); ii) within a clonal bacterial population some phenotypic variants specialise to reduce antibiotic accumulation in multiple ways, from delaying accumulation to reducing accumulation levels. To further test this latter hypothesis, we measured the correlation between different kinetic parameters for each drug data set (Fig. 2 and Table S2). We found a significantly negative correlation between *t*_*0*_ and *k*_*1*_ for the accumulation of polymyxin B, octapeptin and roxithromycin probes; we also found a significantly negative correlation between *t*_*0*_ and *F*_*max*_ for the accumulation of polymyxin B, octapeptin, linezolid and trimethoprim probes and a significantly positive correlation between *k*_*1*_ and *F*_*max*_ for the accumulation of polymyxin B, ciprofloxacin and roxithromycin probes (Fig. 2 and Table S2). Taken together these data suggest that within a clonal population some phenotypic variants specialise to reduce accumulation of a wide range of commonly employed antibiotics.

Furthermore, we also used our mathematical framework to test the hypothesis that treatment with each antibiotic gives rise to a unique accumulation profile that permits identifying and classifying the antibiotic in use, which is important in the context of drug development. Using statistical classification with only two kinetic parameters (*t*_*0*_ and *k*_*1*_, i.e. the two parameters that can be rapidly measured experimentally), we found that treatment with membrane targeting probes is correctly classified against treatment with intracellular targeting probes with 99% accuracy (1,075 cells analysed, Fig. 2D). Moreover, treatment with polymyxin B, tachyplesin or octapeptin was correctly classified among treatments with the other two membrane targeting probes with 77%, 76% and 64%, respectively (Fig. 2E-G). Finally, treatment with linezolid, trimethoprim, ciprofloxacin or roxithromycin was correctly classified among treatments with the other three intracellular targeting probes with 97%, 84%, 64% and 86% accuracy, respectively (Fig. 2H-K). It is worth noting that we obtained similar levels of accuracy when we run such statistical classifications using the full set of kinetic accumulation parameters (i.e. *t*_*0*_, *k*_*1*_, *k*_*2*_, *d*_*r*_ and *d*_*c*_), further demonstrating that measuring only *t*_*0*_ and *k*_*1*_ provides a good description of the antibiotic accumulation process.

Taken together, these data strongly suggest the existence of a unique accumulation pattern for the specific antibiotic in use. Therefore our novel experimental and theoretical framework will enable the classification of novel antibiotic compounds according to their kinetic accumulation parameters. As such this platform could be utilized for rapid phenotyping of bacterial populations ultimately in clinical antibiotic testing.

### Phenotypic variants with reduced antibiotic accumulation survive antibiotic treatment

Next, we hypothesised that phenotypic variants displaying reduced antibiotic accumulation also better survive antibiotic treatment, the correlation between antibiotic uptake and efficacy remaining poorly investigated(17). We decided to focus on the macrolide roxithromycin since a large number of phenotypic variants displayed reduced roxithromycin accumulation (Fig. S2). When we measured the elongation rate of individual cells while they were being dosed with roxithromycin-NBD dissolved in LB, we found two distinct cellular responses. While the majority of cells stopped growing during drug exposure (Fig. 3A), some phenotypic variants within the same clonal *E. coli* population continued elongating for the entire duration of drug treatment (Fig. 3B).

**Figure 3.**
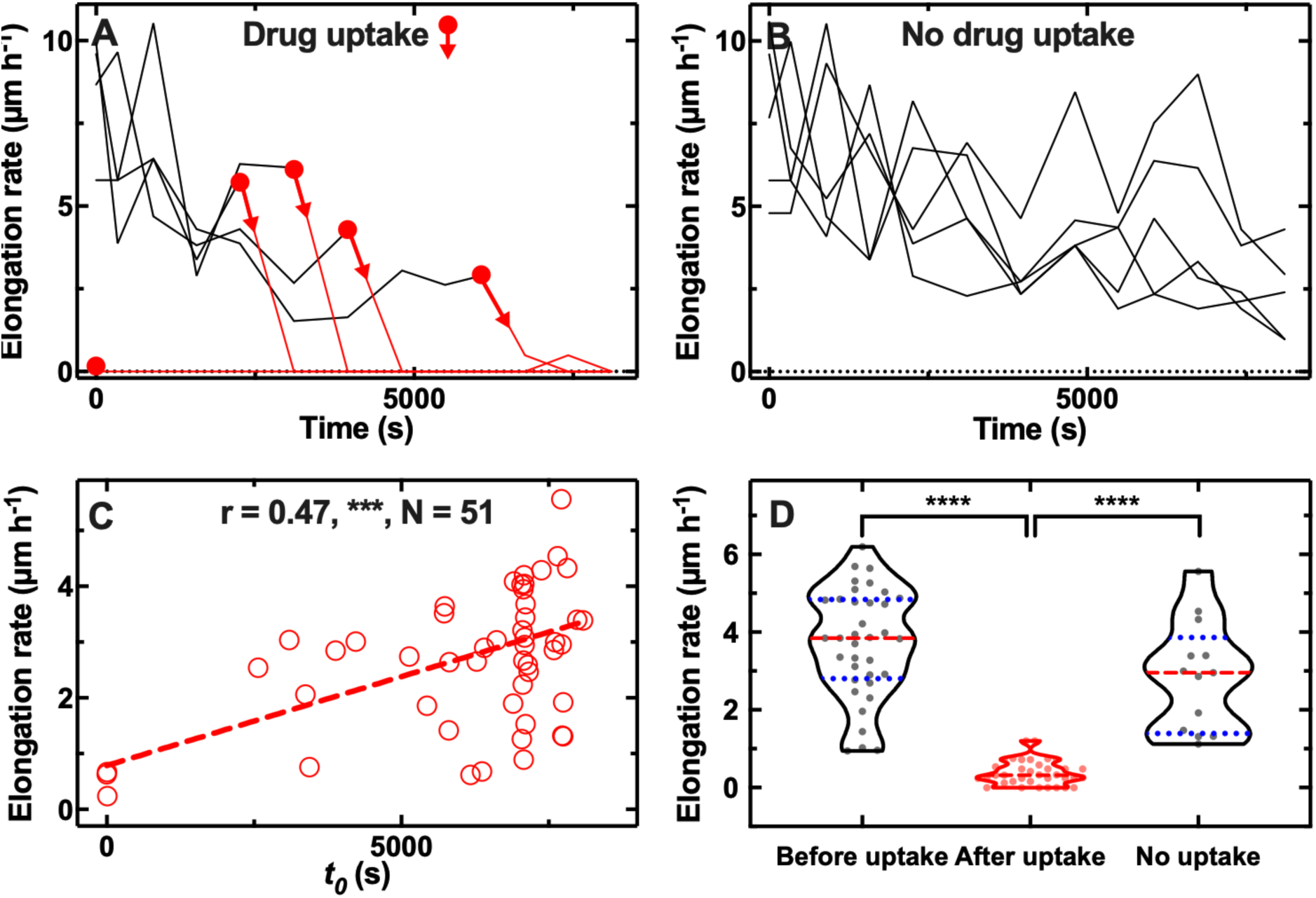
Correlation between antibiotic efficacy and antibiotic accumulation. Temporal patterns of elongation rate during exposure to the fluorescent derivative of roxithromycin for **A)** five representative *E. coli* bacteria that accumulated the drug and **B)** five representative *E. coli* bacteria that did not accumulate the drug. The fluorescent derivative of roxithromycin was delivered at t = 0 at a concentration of 46 μg mL^-1^ and was dissolved in LB, circles and arrows indicate *t*_*0*_, the time point at which each bacterium started to accumulate the drug (i.e. bacterial fluorescence signal became distinguishable from the background). **C)** Correlation between each bacterium *t*_*0*_ and its average elongation rate throughout exposure to the fluorescent derivative of roxithromycin (i.e. 0 < t < 8100 s). r is the Pearson coefficient quantifying the correlation above, ***: p-value < 0.001, N = 52 bacteria. **D)** Average elongation rates for bacteria that had not yet started (before uptake) or had started (after uptake) accumulating the fluorescent derivative of roxithromycin, as well as for bacteria that did not accumulate the drug (no uptake). The red dashed and blue dotted lines within each violin plot represent the median and quartiles of each data set, respectively. Paired t-test between elongation rates before and after onset in accumulation: ****, p-value < 0.0001, N = 36 pairs. Unpaired t-test between the elongation rates of bacteria that did not take up the drug compared to the elongation rate of bacteria that had not yet started taking up the drug: not significant, p-value = 0.07, N = 13 and 36 bacteria, respectively. Unpaired t-test between the elongation rates of bacteria that did not take up the drug compared to the elongation rate of bacteria that had started taking up the drug: ****, p-value < 0.0001, N = 13 and 36 bacteria, respectively.

Furthermore, there were significant cell-to-cell differences in the time at which cells stopped growing (Fig. 3A). Notably, this time coincided with the onset in roxithromycin-NBD accumulation (*t*_*0*_, indicated by circles and arrows in Fig. 3A), whereas phenotypic variants that continued growing did not accumulate roxithromycin-NBD for the entire duration of the treatment (Fig. 3B). These data suggest a strong link between reduced antibiotic accumulation and survival to antibiotic treatment. In fact, we found a strong positive correlation between the onset of roxithromycin-NBD accumulation and the average elongation rate during exposure to roxithromycin-NBD (r = 0.49, ***, Fig. 3C). Moreover, individual bacteria that accumulated roxithromycin-NBD displayed a drastically reduced elongation rate after roxithromycin-NBD accumulation started compared to their elongation rate before uptake (**** paired t-test, Fig. 3D). Phenotypic variants that did not accumulate roxithromycin-NBD instead displayed an elongation rate that was not significantly different compared to the elongation rate of bacteria that had not yet started taking up roxithromycin-NBD (ns unpaired t-test, Fig. 3D). Finally, phenotypic variants that did not accumulate roxithromycin-NBD displayed an elongation rate that was significantly higher compared to the elongation rate of bacteria that had started taking up roxithromycin-NBD (**** unpaired t-test, Fig. 3D).

Taken together these data demonstrate that cell-to-cell differences in drug accumulation have important consequences on the outcome of antibiotic therapy, prompting us to investigate the mechanisms underlying phenotypic variants with delayed or reduced antibiotic accumulation.

### The microcolony architecture affects heterogeneity in antibiotic accumulation

Firstly, we tested the hypothesis that these phenotypic variants reduced antibiotic accumulation because of the presence of other bacteria (i.e. screening cells) between them and the main microfluidic chamber, where the drug is injected. To test this hypothesis, we classified our data in subpopulations of bacteria that had zero, one, two, three or four screening cells between themselves and the main microfluidic chamber (see Inset in Fig. 4E where the drug diffuses from left to right).

**Figure 4.**
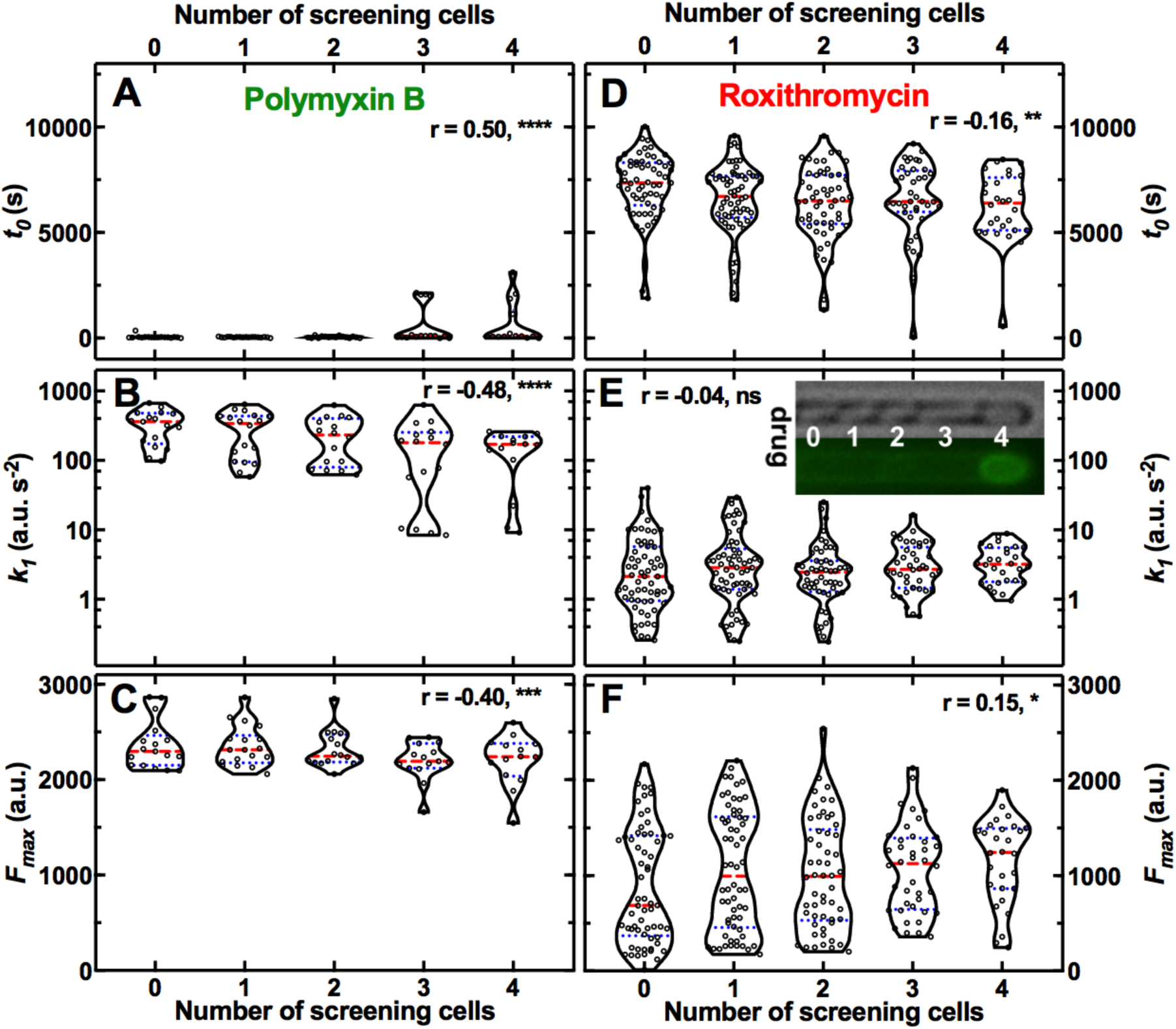
Effect of the presence of screening cells on the accumulation of antibiotics in single cells. Dependence of the kinetic parameters *t*_*0*_, *k*_*1*_, and *F*_*max*_ for the accumulation of fluorescent derivatives of polymyxin B **A-C**) and roxithromycin **D-F**) on the number of screening cells between the bacterium under investigation and the main microfluidic chamber where the drug is continuously injected. Each data point is the value of a kinetic parameter inferred for an individual bacterium from the data in Fig. S1 using our mathematical model, N = 103 and 265 for polymyxin B and roxithromycin, respectively. The red dashed and blue dotted lines within each violin plot represent the median and quartiles of each data set, respectively. r is the Pearson coefficient quantifying the correlation between each inferred kinetic parameter and the number of screening cells in front of each bacterium. ns: not significant correlation, *: p-value < 0.05, **: p-value < 0.01, ***: p-value < 0.001, ****: p-value < 0.0001. Inset: representative brightfield and fluorescence images illustrating, from left to right, a bacterium screened by 0, 1, 2, 3, and 4 cells, respectively; roxithromycin-NBD was injected in the main microfluidic chamber in the left-hand side of the image and diffused from left to right. The fluorescence image shows early roxithromycin-NBD accumulation in the bacterium screened by the highest number of cells.

For polymyxin B we observed that increasing the number of screening cells increased *t*_*0*_ while reducing *k*_*1*_ and *F*_*max*_ (Pearson correlation coefficient r = 0.50, −0.48 and −0.40,****, **** and ***, respectively, Fig. 4A-C). Moreover, octapeptin and tachyplesin displayed strong negative correlation between *k*_*1*_ and the number of screening cells (r = −0.63 and −0.67, respectively, ****); octapeptin also displayed a strong positive correlation between *t*_*0*_ and the number of screens (r = 0.71, ****). These data were in accordance with our hypothesis that screening cells transiently decrease the pool of drug molecules available for screened cells until the bacteria closer to the main chamber reach antibiotic accumulation saturation levels. These data provide a mechanistic understanding for the large heterogeneity in *t*_*0*_ measured for such membrane-targeting probes (Fig. S11). In contrast with our hypothesis, for roxithromycin we found that increasing the number of screens in front of a cell reduced *t*_*0*_ and increased *F*_*max*_ (r = −0.16 and 0.15, ** and *, respectively, Fig. 4D-F). Moreover, both ciprofloxacin and linezolid displayed a strong negative correlation between *t*_*0*_ and the number of screens (r = −0.53 and −0.28, **** and ***, respectively); ciprofloxacin also displayed a strong positive correlation between *k*_*1*_ and the number of screens (r = 0.32, ***). These novel findings were unexpected and were not dictated by oxygen limitation or low metabolic activity as in the case of biofilms(64). In fact, we(47) and others(65) have previously demonstrated that nutrients, including oxygen and metabolites, uniformly distribute across the whole length of bacteria hosting channels in our microfluidic device.

Taken together these findings suggest non-trivial and drug-specific effects of the bacterial microcolony architecture on the dynamics of drug accumulation in individual bacteria, a novel phenotypic feature that should be taken into account when designing and optimising new drugs and therapies. Moreover, mechanisms other than the microcolony architecture must underlie phenotypic variants with reduced antibiotic accumulation. In fact, we registered significant cell-cell differences in antibiotic accumulation even within the same subpopulation of bacteria with the same number of screening cells; these differences were more pronounced for antibiotic with intracellular targets compared to membrane targeting antibiotics (e.g. roxithromycin and polymyxin B, respectively, in Fig. 4)

### Cell-to-cell differences in growth rate before treatment underlie heterogeneity in antibiotic accumulation

In order to further dissect the mechanisms underlying phenotypic variants with reduced antibiotic accumulation, we took advantage of continuous live-cell imaging to track individual bacteria for the two-hour growth period in LB before incubation in each antibiotic. This permitted us to investigate links between each bacterium’s growth and its capability to avoid or delay antibiotic accumulation. We investigated the correlation between elongation rate before treatment and the kinetic parameters describing the accumulation of two representative membrane-targeting antibiotics, i.e. octapeptin and tachyplesin, and two representative antibiotics with intracellular targets, i.e. trimethoprim and roxithromycin.

We did not find any significant correlation between single-cell elongation rate before treatment and any of the kinetic parameters describing the accumulation of octapeptin and trimethoprim (Fig. S13A-C and S13G-I, respectively). However, we found a positive correlation between single-cell elongation rate before treatment and *k*_*1*_ for tachyplesin (r = 0.59, **, Fig. S13E), suggesting that the latter accumulated faster in fast growing cells. On the contrary, for roxithromycin, we found a significantly positive correlation between single-cell elongation rate before treatment and *t*_*0*_ and a significantly negative correlation between single-cell elongation rate before treatment and *F*_*max*_ (r = 0.66 and −0.54, **** and ***, respectively, Fig. 5A and 5C), but no correlation with cell size (Fig. S4). Furthermore, we found that, as expected, the average elongation rate significantly decreased after roxithromycin-NBD addition (5.2 ± 3.7 μm h^-1^ vs 3.7 ± 2.3 μm h^-1^, before and after drug addition, respectively, ****, Fig. S14A). Moreover, we also found a significantly positive correlation between single-cell elongation rate before treatment and single-cell elongation rate during treatment (r = 0.34, *, Fig. S14A). Finally, to further verify that these findings were not due to drug labelling, we performed these experiments with unlabelled roxithromycin confirming a significantly positive correlation between single-cell elongation rate before treatment and single-cell elongation rate during treatment (r = 0.47, ***, Fig. S14B).

**Figure 5.**
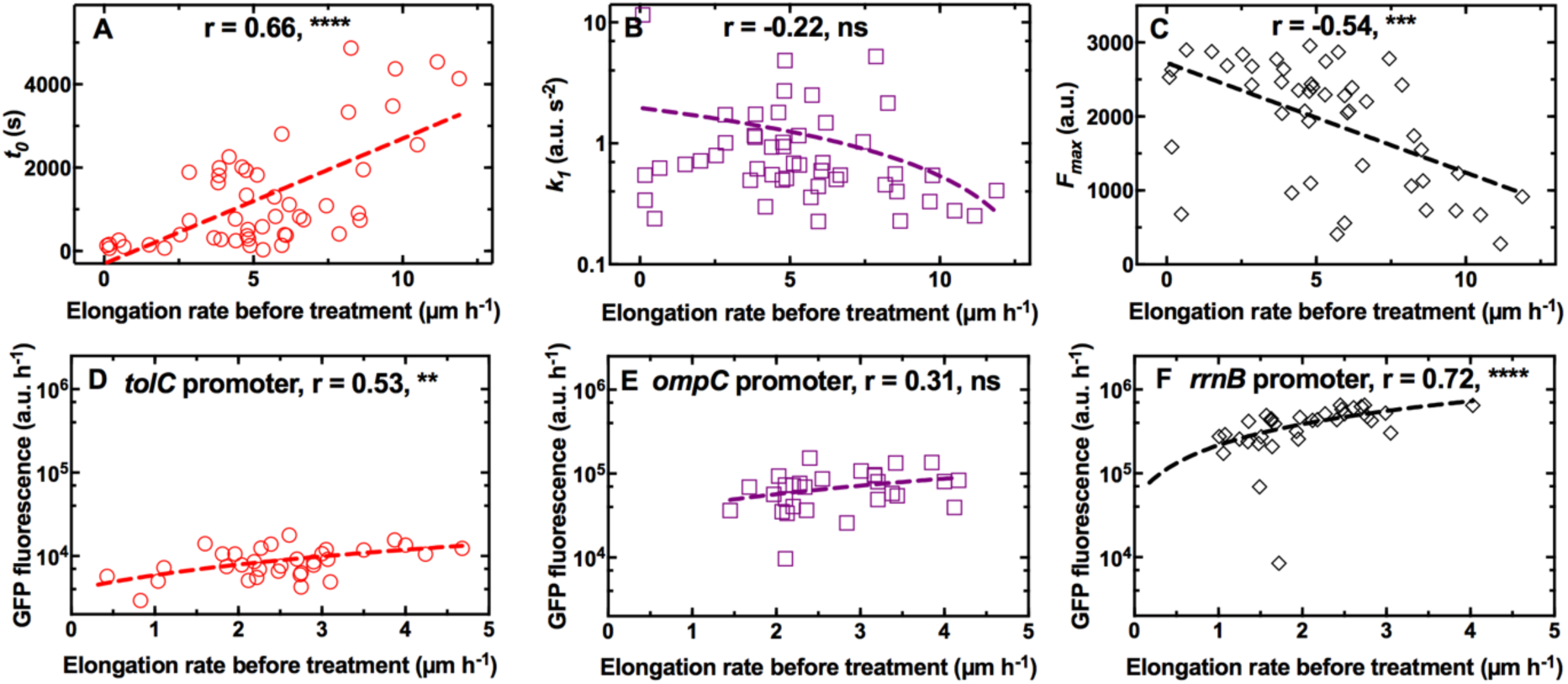
Differential cell growth and expression of key molecular pathways underlie heterogeneity in roxithromycin accumulation. **A-C**) Correlation between the single-cell kinetic parameters *t*_*0*_, *k*_*1*_ and *F*_*max*_ describing the accumulation of roxithromycin-NBD and the bacterial elongation rate during the two-hour growth period preceding antibiotic treatment (see Methods). Measurements were carried out on N = 50 individual *E. coli*, collated from biological triplicate, before and after exposure to 192 μg mL^-1^ roxithromycin-NBD dissolved in M9. **D-F**) Correlation between the single-cell GFP fluorescence as a proxy for the expression of *tolC, ompC* and *rrnB* promoters and the bacterial elongation rate during the two hour growth period preceding antibiotic treatment (see Methods). r is the Pearson coefficient quantifying the correlation between each inferred kinetic parameter and the corresponding elongation rate of each cell. ns: not significant correlation, **: p-value < 0.01, ***: p-value < 0.001, ****: p-value < 0.0001. Dashed lines are linear regressions to the data. Measurements were carried out on N = 34, 30 and 35 individual *E. coli* collated from biological triplicate for the *tolC, ompC* and *rrnB* reporter strains, respectively.

These data demonstrate that phenotypic variants displaying reduced roxithromycin accumulation are fast growing bacteria that also better survive roxithromycin treatment, thus establishing, for the first time, a strong link between heterogeneity in antibiotic efficacy and cell-to-cell differences in antibiotic accumulation. These novel findings are surprising considering that phenotypic survival to antibiotics has traditionally been linked to slow growth, low metabolic activity and bacterial dormancy(66–68). In contrast, here we show that fast growth facilitates delayed roxithromycin accumulation as well as reducing the amount of macrolide accumulating in individual bacteria at steady state this decreasing roxithromycin efficacy.

### Single-cell ribosome and efflux pump abundance underlies heterogeneity in macrolide accumulation

In order to determine the molecular mechanisms underpinning phenotypic variants with reduced roxithromycin accumulation, we investigated whether heterogeneity in bacterial growth rate could be linked to heterogeneity in the expression of key molecular pathways underlying roxithromycin accumulation. We hypothesised that heterogeneity in *t*_*0*_ could be linked to cell-to-cell differences in the capability to pump antibiotics out from the cell, thus delaying the onset of accumulation. *tolC*, which encodes the outer membrane channel of the multi-drug efflux pump AcrAB-TolC and the macrolide efflux pump MacAB-TolC(17), was the most highly expressed efflux pump related gene according to our transcriptomic data of *E. coli* cultures growing on LB for a period of two hours after dilution of an overnight culture (Table S3 and(69)). Therefore, we used a *tolC* transcriptional reporter strain(8) to establish a link between the kinetic parameter *t*_*0*_, single-cell elongation rate and *tolC* expression during the two hour growth period before exposure to roxithromycin. In line with our hypothesis above, we found a positive correlation between the expression of *tolC* and single-cell elongation rate during the two-hour growth period before exposure to roxithromycin (r = 0.53, **, Fig. 5D).

Next, we hypothesised that heterogeneity in the rate of drug uptake *k*_*1*_ could be ascribed to cell-to-cell differences in the expression of outer membrane porins allowing antibiotic passage across the outer membrane. *ompC*, which encodes the outer membrane protein OmpC facilitating influx of several antibiotics(17,70), was the most highly expressed outer membrane protein encoding gene according to our transcriptomic data at the population level (Table S3 and(69)). In contrast with our hypothesis, we did not find a significant correlation between *ompC* expression and single-cell elongation rate during the two-hour growth period before drug exposure (r = 0.31, ns, Fig. 5E). These data demonstrate that bacteria growing at different rates do not display significantly different levels of *ompC* expression and accordingly we did not find significant correlation between cell growth and *k*_*1*_ (Fig. 5B).

Finally, we hypothesised that cell growth and saturation levels in roxithromycin accumulation could depend on the ribosomal content (i.e. the drug target) at the single-cell level. Accordingly, we found a strong positive correlation between the expression of the ribosomal promoter *rrnB* and single-cell elongation rate during the two-hour growth period before exposure to roxithromycin (r = 0.72, ****, Fig. 5F).

Taken together these data shed light on the molecular mechanisms underpinning the observed heterogeneity in the intracellular accumulation of the macrolide roxithromycin: fast growing variants reduce the intracellular accumulation of roxithromycin, and thus better survive treatment with this drug, via elevated ribosomal content and, to a lesser extent, higher expression of efflux pumps. These data call into question the current consensus that metabolically inactive or dormant bacteria better survive antibiotic challenge(66– 68,71).

### External manipulation of the heterogeneity in antibiotic accumulation

Building on the molecular understanding gained above, we then set out to establish whether phenotypic variants displaying reduced roxithromycin accumulation could be suppressed either genetically or chemically. In order to do so, we employed a *ΔtolC* knock-out mutant and found that, when investigating roxithromycin accumulation, *t*_*0*_ was significantly lower and *k*_*1*_ was significantly higher in the *ΔtolC* mutant compared to the parental strain (Fig. 6A and 6B, respectively).

**Figure 6.**
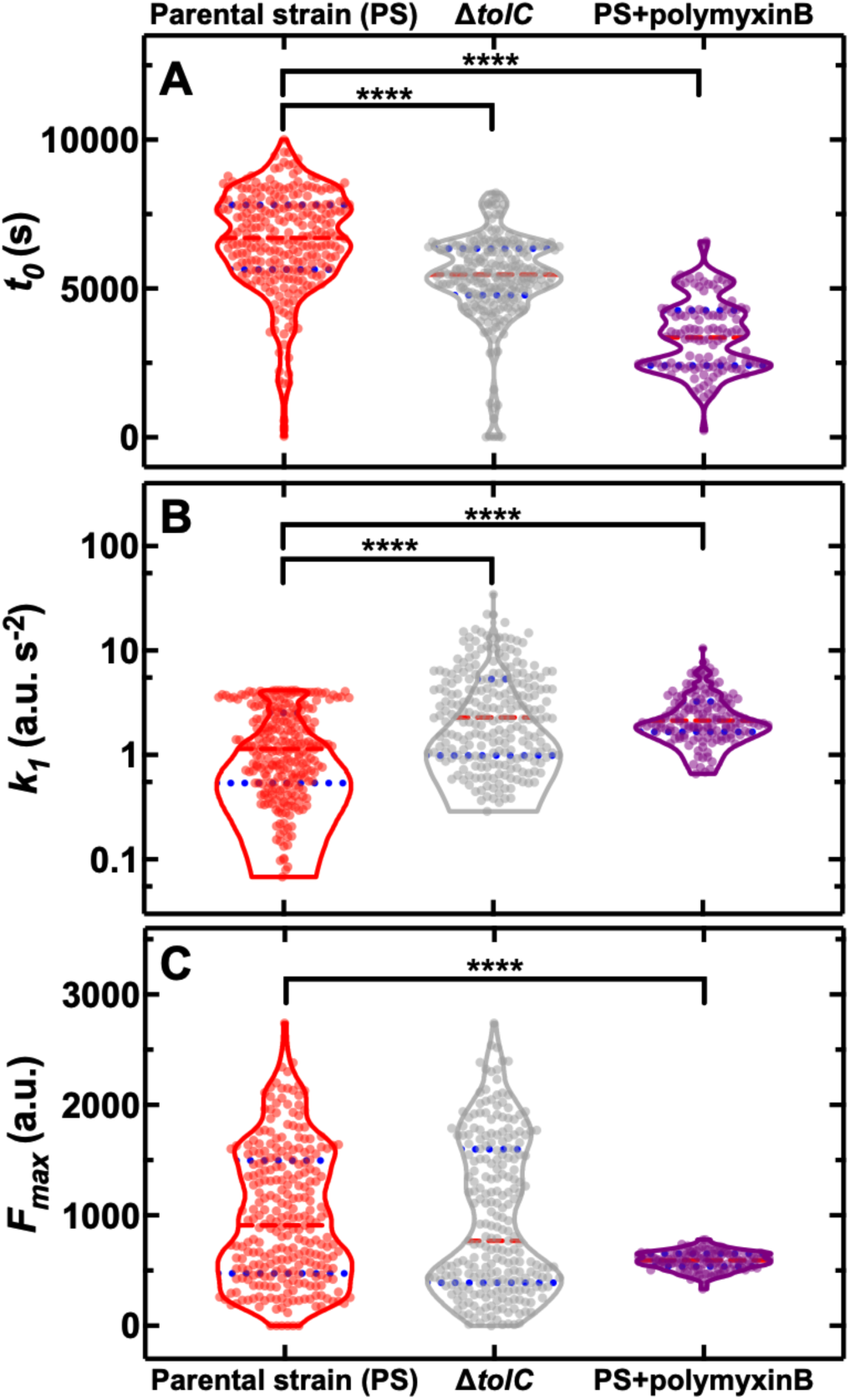
Genetic and chemical manipulation of heterogeneity in drug accumulation. Distributions of single-cell values for the kinetic parameters **A**) *t*_*0*_, **B**) *k*_*1*_ and **C**) *F*_*max*_ describing the accumulation of the fluorescent derivative of roxithromycin (at 46 μg mL^-1^ in M9) in the *E. coli* BW25113 parental strain (PS), the knock-out mutant *ΔtolC* and the parental strain co-treated with unlabelled polymyxin B at 1 μg mL^-1^ extracellular concentration. The red dashed and blue dotted lines within each violin plot represent the median and quartiles of each data set, respectively. ****: p-value<0.0001. N = 262, 241 and 116 individual parental strain *E. coli* treated with the roxithromycin probe, *ΔtolC E. coli* treated with the roxithromycin probe and parental strain *E. coli* co-treated with the roxithromycin probe and 1 μg mL^-1^ unlabelled polymyxin B.

However, we also found *ΔtolC* phenotypic variants with reduced roxithromycin accumulation and even higher levels of heterogeneity in the three kinetic parameters for the *ΔtolC* mutant compared to the parental strain (CV of 27% vs 25%, 114% vs 80%, 72% vs 62% for *t*_*0*_, *k*_*1*_, and *F*_*max*_, respectively, Fig. 6). These data demonstrate that targeting efflux might not be a promising avenue to reduce heterogeneity in drug accumulation and confirm that the observed heterogeneity in roxithromycin accumulation is not exclusively underpinned by cell-to-cell differences in efflux pump expression. Since we demonstrated that heterogeneity in porin expression does not underpin cell-to-cell differences in roxithromycin accumulation, we hypothesised that the composition and permeability of the lipid bilayer making up the bacterial outer membrane could underlie heterogeneity in roxithromycin accumulation. If this were true, the heterogeneity in roxithromycin accumulation could be chemically manipulated by using agents that permeabilise the outer membrane, such as polymyxin B(72). Accordingly, when we treated the parental strain with roxithromycin-NBD at 46 μg mL^-1^ in combination with unlabelled polymyxin B at 1 μg mL^-1^ extracellular concentration, we found a significant decrease in the heterogeneity of *k*_*1*_ and *F*_*max*_ compared to roxithromycin-NBD treatment alone (CV of 59% vs 80%, 14% vs 62%, respectively, Fig. 6B and 6C). Additionally, the accumulation dynamics of roxithromycin-NBD in the presence of unlabelled polymyxin B was significantly earlier and faster compared to that measured in the absence of polymyxin B (Fig. 6). Taken together, these data suggest that phenotypic variants displaying reduced roxithromycin accumulation might have a significantly more impermeable outer membrane than phenotypically susceptible bacteria, possibly due to differences in lipid composition and packing and that targeting the outer membrane might be a viable avenue for suppressing variants with reduced intracellular antibiotic accumulation.

## Discussion

Bacterial slow growth has often been associated with decreased antibiotic susceptibility(66,73,74) with few exceptions(75,76). Moreover, a recent paper suggested that phenotypic variants accumulate lower levels of phenoxymethylpenicillin while being in a dormant state before treatment(59). In striking contrast, here we provide compelling evidence that fast growth and elevated ribosomal content better prepare phenotypic variants for avoiding the intracellular accumulation of macrolides, a finding that needs to be considered when designing antibiotic therapy.

A linear correlation between ribosomal abundance and growth rate has previously been found via ensemble measurements obtained on exponentially growing *E. coli* supplied with nutrients of increasing quality in the absence of antibiotics(77). Our findings enrich the current understanding of the interdependence of cell growth and ribosomal content demonstrating that this correlation holds within an isogenic population homogeneously exposed to the same medium.

Previous ensemble measurements have demonstrated that fast growth on high quality nutrients decreases *E. coli* growth inhibition by antibiotics that irreversibly bind to ribosomes (such as roxithromycin (78)) compared to slower growth on poor quality nutrients(79). Here, we offer a mechanistic understanding of this unexpected finding, showing that reduced growth inhibition in fast growing cells is dictated by growth-dependent transport rates, as fast growing variants displayed reduced macrolide accumulation. Importantly, we demonstrated that this phenotypic response is found not only at the population-level(79), but also within an isogenic population.

These new data can be rationalised by considering that in fast growing variants a fraction of leading actively translating ribosomes(80) escapes roxithromycin binding, while other ribosomes stall after accumulating roxithromycin. Drug-free active ribosomes continue to facilitate essential cellular processes including efflux that can reduce macrolide accumulation. Accordingly, we found that before antibiotic treatment fast growing variants also displayed a significantly higher expression of the efflux promoter *tolC* compared to slow growing cells. Moreover, the deletion knockout *ΔtolC* displayed significantly earlier and faster accumulation of roxithromycin compared to the parental strain, confirming that roxithromycin is a substrate of the AcrAB- and MacAB-TolC efflux pumps(24). However, this mutant exhibited accumulation heterogeneity levels comparable to the parental strain. These data suggest that phenotypic variants reduce antibiotic accumulation using processes other than efflux alone, in contrast with previous findings(59), and in accordance with our data on the key role played by heterogeneity in ribosomal abundance.

Our data also revealed a strong correlation between the accumulation of roxithromycin and the effect of this antibiotic on cell growth down to the scale of the individual cell. This suggests that phenotypic variants with reduced antibiotic accumulation could be an important factor contributing to phenotypic resistance to antibiotics(2,8,73,81). This fundamentally new knowledge calls for a major rethink about phenotypic resistance to antibiotics that is currently centred around target deactivation or modification(73,82,83) with very little known about the correlation between antibiotic accumulation and antibiotic efficacy(17,59).

Experimental evidence suggests that both macrolides and polymyxins use the self-promoted uptake pathway. Moreover, polymyxins have a higher affinity to the LPS compared to macrolides and increase the permeability of the outer membrane to other freely diffusing antibiotic molecules (23). Accordingly, we observed that growth-dependent transport rates were not dictated by heterogeneity in the expression of OmpC, which is a major route of antibiotic influx via the hydrophilic pathway(84). Our data show instead that the phenotypic variants that avoid roxithromycin accumulation can be suppressed by delivering roxithromycin in combination with polymyxin B. Moreover, roxithromycin accumulated at lower saturation levels in the presence of polymyxin B as expected due to competitive binding to the LPS.

These data suggest that heterogeneity in roxithromycin accumulation could also be due to cell-to-cell differences in LPS composition. It is conceivable that phenotypic variants within the clonal population might have a decreased ethanolamine content. This would result in an increased negative charge of the LPS core and a decreased permeability to roxithromycin but not to polymyxin B(85) in accordance with our data. It is also conceivable that phenotypic variants within the clonal population might display esterification of the core-lipid A phosphates(63). However, this would result in decreased permeability to both roxithromycin and polymyxin B in contrast with our data showing i) comparatively smaller cell-to-cell differences in polymyxin B accumulation (beyond the heterogeneity generated by the microcolony architecture) and ii) that adding polymyxin B suppresses the heterogeneity in roxithromycin accumulation. Finally, it has been suggested that macrolides use the hydrophobic pathway (86). It is conceivable that phenotypic variants within the clonal population might display a higher expression of *lpxA* and thus reduced permeability to roxithromycin; however, this hypothesis remains to be tested.

We further demonstrate that the presence of phenotypic variants that avoid antibiotic accumulation is not dictated by the microcolony architecture (as represented by bacterial cell position within a microfluidic channel). However, our data offer a mechanistic understanding of previous work in clinical settings suggesting that macrolides, quinolones, and oxazolidinones are more effective within infecting biofilms compared to glycopeptides and polymyxins(64,87). In fact, we demonstrate that antibiotics with intracellular targets accumulate more readily and to higher saturation levels in bacteria within the inner core of the colony. In contrast, membrane targeting drugs accumulate more readily, faster and at higher saturation levels in bacteria at the outer rim of the colony. This drug-specific effect of colony architecture on drug accumulation must rely on growth- and efflux-independent mechanisms. In fact, we did not find significant correlations between the position of a cell within the colony and neither the expression of *tolC, ompC* or *rrnB* nor the bacterial elongation rate (p-value = 0.13, 0.13, 0.46 and 0.34, respectively).

In conclusion, this work reveals hitherto unrecognised phenotypic variants that avoid antibiotic accumulation within bacterial populations. In contrast with the current consensus, we demonstrate that fast growing phenotypic variants avoid macrolide accumulation and survive treatment due to elevated ribosomal content. We show that it is possible to phenotypically engineer clonal bacterial populations by eradicating phenotypic variants currently avoiding antibiotic accumulation. These data give strength to recent evidence that administered doses of polymyxins can be lowered in combination therapies(40) and demonstrating that roxithromycin could be repurposed against gram-negative bacteria. Finally, our novel single-cell approach reveals that each antibiotic is characterised by a unique accumulation pattern and thus will in future allow to classify new leading antibiotic compounds(88–91) using their kinetic accumulation parameters, guiding medicinal chemistry(24) whilst avoiding biases previously introduced by activity-dependent screenings(31).

## Materials and Methods

### Chemicals and cell culture

All chemicals were purchased from Fisher Scientific or Sigma-Aldrich unless otherwise stated. Lysogeny broth (LB) medium (10 g L^-1^ tryptone, 5 g L^-1^yeast extract, and 0.5 g L^-1^ NaCl) and LB agar plates (LB with 15 g L^-1^ agar) were used for planktonic growth and setting up overnight cultures. Glucose-free M9-minimal media, used to dissolve fluorescent antibiotic derivatives was prepared using 5× M9 minimal salts (Merck), diluted as appropriate, with additional 2 mM MgSO_4_, 0.1 mM CaCl_2_, 3 μM thiamine HCl in Milli-Q water. Stock solutions of polymyxin B, octapeptin, tachyplesin, vancomycin, linezolid, roxithromycin and trimethoprim were obtained by dissolving these compounds in dimethyl sulfoxide; ciprofloxacin instead was dissolved in 0.1 M HCl in Milli-Q water. These stock solutions were prepared at a concentration of 640 μg mL^-1^. *Escherichia coli* BW25113 was purchased from Dharmacon (GE Healthcare). *ompC, tolC* and *rrnB* reporter strains of an *E. coli* K12 MG1655 promoter library(92) were purchased from Dharmacon (GE Healthcare). Plasmids were extracted and transformed into chemically competent *E. coli* BW25113 as previously reported(93). *Staphylococcus aureus* ATCC 25923, *Pseudomonas aeruginosa* PA14 flgK::Tn5(Tcr) (the deletion of the flagellum FlgK facilitated holding cells in the hosting channel thanks to the reduced bacterial motility) and *Burkholderia cenocepacia* K56-2 were kindly provided by A. Brown and S. van Houte. All strains were stored in 50% glycerol stock at −80 °C. Streak plates for each strain were produced by thawing a small aliquot of the corresponding glycerol stock every 2 weeks and plated onto LB agar. Overnight cultures were prepared by picking a single bacterial colony from a streak plate and growing it in 100 mL fresh LB medium on a shaking platform at 200 rpm and 37 °C for 17 h.

### Synthesis of fluorescent derivatives of antibiotics

Fluorescent antibiotic derivatives from trimethoprim(56) (antifolate), linezolid(53) (oxazolidinone), ciprofloxacin(55) (fluoroquinolone) and roxithromycin(52) (macrolide) were prepared as previously described. Vancomycin(94) (glycopeptide), polymyxin(95) and octapeptin(96) (both lipopeptides) and tachyplesin(97) (antimicrobial peptide) analogues were designed and synthesised based on structure-activity-relationship studies and synthetic protocols reported in prior publications, introducing an azidolysine residue for the subsequent ‘click’ reactions with nitrobenzoxadiazole (NBD)-alkyne. Additionally, a fluorescent derivative of roxithromycin using the fluorophore dimethylamino-coumarin-4-acetate (DMACA) was synthesised and used only to determine the impact of labelling on single-cell antibiotic accumulation.

### Determination of minimum inhibitory concentration

Single colonies of *E. coli* BW25113 were picked and cultured overnight in cation-adjusted Mueller Hinton broth (CAMHB) at 37 °C, then diluted 40-fold and grown to OD_600_ = 0.5. 60 μL of each antibiotic or fluorescent antibiotic derivative stocks were added to the first column of a 96-well plate. 40 μL CAMHB was added to the first column, and 30 μL to all other wells. 70 μL solution was then withdrawn from the first column and serially transferred to the next column until 70 μL solution withdrawn from the last column was discharged. The mid-log phase cultures (i.e. OD_600_ = 0.5) were diluted to 10^6^ colony forming units (c.f.u.) ml^-1^ and 30 μL was added to each well, to give a final concentration of 5×10^5^ c.f.u. ml^-1^. Each plate contained two rows of 12 positive control experiments (i.e. bacteria growing in CAMHB without antibiotics) and two rows of 12 negative control experiments (i.e. CAMHB only). Plates were covered with aluminium foil and incubated at 37 °C overnight. The minimum inhibitory concentrations (MICs) of fluorescent derivatives of polymyxin B, octapeptin, tachyplesin, vancomycin, linezolid, roxithromycin, ciprofloxacin, trimethoprim and each corresponding parental antibiotic against *E. coli* BW25113 were determined visually, with the MIC being the lowest concentration well with no visible growth (compared to the positive control experiments).

### Fabrication of the microfluidic devices

The mould for the mother machine microfluidic device(65) was obtained by pouring epoxy onto a polydimethylsiloxane (PDMS, Dow Corning) replica of the original mould containing 12 independent microfluidic chips (kindly provided by S. Jun). Each of these chips is equipped with approximately 6000 lateral microfluidic channels with width and height of 1 μm each and a length of 25 μm. These lateral channels are connected to a main microfluidic chamber that is 25 μm and 100 μm in height and width, respectively. PDMS replicas of this device were realised as previously described(98). Briefly, a 10:1 (base:curing agent) PDMS mixture was cast on the mould and cured at 70 °C for 120 min in an oven. The cured PDMS was peeled from the epoxy mould and fluidic accesses were created by using a 0.75 mm biopsy punch (Harris Uni-Core, WPI). The PDMS chip was irreversibly sealed on a glass coverslip by exposing both surfaces to oxygen plasma treatment (10 s exposure to 30 W plasma power, Plasma etcher, Diener, Royal Oak, MI, USA). This treatment temporarily rendered the PDMS and glass hydrophilic, so immediately after bonding the chip was filled with 2 μL of a 50 mg/mL bovine serum albumin solution and incubated at 37 °C for 30 min, thus passivating the internal surfaces of the device and preventing subsequent cell adhesion. We have also made available a step-by-step experimental protocol for the fabrication and handling of microfluidic devices for investigating the interactions between antibiotics and individual bacteria(99).

### Imaging single-cell drug accumulation dynamics

An overnight culture was prepared as described above and typically displayed an optical density at 595 nm (OD_595_) around 5. A 50 mL aliquot of the overnight culture above was centrifuged for 5 min at 4000 rpm and 37 °C. The supernatant was filtered twice (Medical Millex-GS Filter, 0.22 μm, Millipore Corp.) to remove bacterial debris from the solution and used to resuspend the bacteria in their spent LB to an OD_600_ of 75. A 2 μL aliquot of this suspension was injected in the microfluidic device above described and incubated at 37 °C. The high bacterial concentration favours bacteria entering the narrow lateral channels from the main microchamber of the mother machine(8). We found that an incubation time between 5 and 20 min allowed filling of the lateral channels with, typically, between one and three bacteria per channel. Shorter incubation times were required for motile or small bacteria, such as *P. aeruginosa* and *S. aureus*, respectively. An average of 80% of lateral channels of the mother machine device were filled with bacteria. The microfluidic device was completed by the integration of fluorinated ethylene propylene tubing (1/32” × 0.008”). The inlet tubing was connected to the inlet reservoir which was connected to a computerised pressure-based flow control system (MFCS-4C, Fluigent). This instrumentation was controlled by MAESFLO software (Fluigent). At the end of the 20 min incubation period, the chip was mounted on an inverted microscope (IX73 Olympus, Tokyo, Japan) and the bacteria remaining in the main microchamber of the mother machine were washed into the outlet tubing and into the waste reservoir by flowing LB at 300 μL h^-1^ for 8 min and then at 100 μL h^-1^ for 2 h. Bright-field images were acquired every 20 min during this 2 h period of growth in LB. Images were collected via a 60×, 1.2 N.A. objective (UPLSAPO60XW, Olympus) and a sCMOS camera (Zyla 4.2, Andor, Belfast, UK). The region of interest of the camera was adjusted to visualise 23 lateral channels per image and images of 10 different areas of the microfluidic device were acquired at each time point in order to collect data from at least 100 individual bacteria per experiment. The device was moved by two automated stages (M-545.USC and P-545.3C7, Physik Instrumente, Karlsruhe, Germany, for coarse and fine movements, respectively). After this initial 2 h growth period in LB, the microfluidic environment was changed by flowing minimal medium M9 (unless otherwise stated) with each of the NBD (unless otherwise stated) fluorescent antibiotic derivatives at a concentration of 46 μg mL^-1^ (unless otherwise stated, also unlabelled ciprofloxacin was delivered at 200 μg mL^-1^) at 300 μL h^-1^ for 8 min and then at 100 μL h^-1^ for 4 h. During this 4 h period of exposure to the fluorescent antibiotic derivative in use, upon acquiring each bright-field image the microscope was switched to fluorescent mode and FITC filter using a custom built Labview software. A fluorescence image was acquired by exposing the bacteria to the blue excitation band of a broad-spectrum LED (CoolLED pE300white, maximal power = 200 mW Andover, UK) at 20% of its intensity (with a power associated with the beam light of 7.93 mW at the sample plane). In the case of unlabelled ciprofloxacin the UV excitation band of such LED was used at 100% of its intensity. These parameters were adjusted in order to maximise the signal to noise ratio. Bright-field and fluorescence imaging during this period was carried out every 5 min. The entire assay was carried out at 37 °C in an environmental chamber (Solent Scientific, Portsmouth, UK) surrounding the microscope and microfluidics equipment.

### Image and data analysis

Images were processed using ImageJ software as previously described(47,48,100), tracking each individual bacterium throughout the initial 2 h period of growth and the following 4 h period treatment with each fluorescent antibiotic derivative. Briefly, during the initial 2 h growth in LB, a rectangle was drawn around each bacterium in each bright-field image at every time point, obtaining its width, length and relative position in the hosting microfluidic channel. Each bacterium’s average elongation rate was calculated as the average of the ratios of the differences in bacterial length over the lapse of time between two consecutive time points. During the following 4 h incubation in the presence of the fluorescent antibiotic derivative, a rectangle was drawn around each bacterium in each bright-field image at every time point obtaining its width, length and relative position in the hosting microfluidic channel. The same rectangle was then used in the corresponding fluorescence image to measure the mean fluorescence intensity for each bacterium that is the total fluorescence of the bacterium normalised by cell size (i.e. the area covered by each bacterium in our 2D images), to account for variations in antibiotic accumulation due to the cell cycle(60). The same rectangle was then moved to the closest microfluidic channel that did not host any bacteria in order to measure the background fluorescence due to the presence of extracellular fluorescent antibiotic derivative in the media. This mean background fluorescence value was subtracted from the bacterium’s fluorescence value. Background subtracted values smaller than 20 a.u. were set to zero since this was the typical noise value in our background measurements. All data were then analysed and plotted using GraphPad Prism 8. Statistical significance was tested using either paired or unpaired, two-tailed, Welch’s *t*-test. Pearson correlation, means, standard deviations, coefficients of variation and medians were also calculated using GraphPad Prism 8.

### Inferring single-cell kinetic parameters of antibiotic accumulation via mathematical modelling

We constructed a minimal model of antibiotic accumulation in order to infer key kinetic parameters quantifying the accumulation of each antibiotic. We modelled antibiotic accumulation using the following set of ordinary differential equations (ODEs):

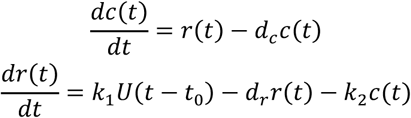

where *U*(*t* – *t*_0_)represents the dimensionless step function:

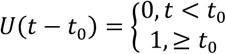

Variable *c*(*t*)represents the intracellular antibiotic concentration (in arbitrary units [a.u.] of fluorescence levels), and *r*(*t*)[a.u. s^-1^] describes the antibiotic uptake rate. With the first equation we described how antibiotic accumulation, *c*(*t*), changes over time as a result of two processes: (i) drug-uptake, which proceeds at a time-varying rate, *r*(*t*); and (ii) drug loss (efflux or antibiotic transformation), which we modelled as a first order reaction with rate constant *d*_*c*_[s^-1^]. With the second equation we described the dynamics of time-varying antibiotic uptake rate, *r*(*t*). The uptake rate starts increasing with a characteristic time-delay (parameter *t*_0_), parameter *k*_1_[a.u. s^-2^] is the associated rate constant of this increase. We also assumed a linear dampening effect (with associated rate constant *d*_*r*_[s^-1^]) to constrain the increase in uptake rate, which allowed us to recapitulate the measured saturation in antibiotic accumulation. In this model the maximum saturation is given by 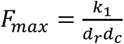. Finally, we introduced an adaptive inhibitory term (rate constant *k*_2_[a.u. s^-2^]) to describe the dip we observed in some single-cell trajectories in Fig. S1 and S2 which we assumed is due to the fact that the presence of drugs intracellularly inhibits further drug uptake. We note that in this model we did not make any a priori assumptions about the mechanisms underlying antibiotic accumulation but rather aimed to capture the dynamics of the measured accumulation data.

Model parameters were inferred from single-cell fluorescence time-traces (see Image and data analysis section) using the probabilistic programming language Stan through its python interface pystan(101). Stan provides full Bayesian parameter inference for continuous-variable models using the No-U-Turn sampler, a variant of the Hamiltonian Monte Carlo method. All No-U-Turn parameters were set to default values except parameter adapt_delta which was set to 0.999 to avoid divergent runs of the algorithm. For each single-cell fluorescence time-trace the algorithm produced 4 chains, each one consisting of 3000 warmup iterations followed by 1000 sampling iterations, giving in total 4000 samples from the parameters’ posterior distribution. For each parameter, the median of the sampled posterior is used for subsequent analysis. For parameter inference, model time was rescaled by the length of the time-trace T, i.e. 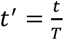 so that time runs between 0 and 1, and model parameters were re-parameterised (and made dimensionless) according to the rules 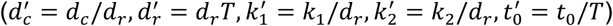. The following diffuse priors were used for the dimensionless parameters, where *U*(*a, b*)denotes the uniform distribution in the range [*a,b*]: 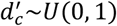 so that uptake rate dynamics are always faster than drug-accumulation dynamics, i.e., 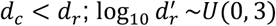 constraining the timescale associated with *d*_*r*_ to be shorter than the timescale of the experiment, i.e., 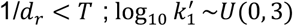 and 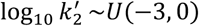, so that the parameter controlling adaptive inhibition is small enough and there is no oscillatory behaviour in the model i.e, 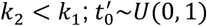,since the transformed time *t*^+^ runs from 0 to 1.

### Statistical classification of the accumulation of antibiotics

For each cell, the marginal posterior distributions of all model parameters (*t*_*0*_, *k*_*1*,_ *k*_*2*_, *d*_*r*_, *d*_*c*_) were summarised using the corresponding first (Q_1_), second (Q_2_) and third (Q_3_) quantiles. For each classification task, a statistical model (classification decision tree) was developed for predicting the drug class for each cell using the summarised parameter posterior distributions as input. Depending on the classification task, either all 5 parameters were considered (5×3=15 predictors) or just parameters *t*_*0*_ and *k*_*1*_ (2×3=6 predictors). Statistical classification was performed using Matlab (method *fitctree*) and the results presented were obtained using 10-fold cross-validation.

## Acknowledgments

U.L. was supported through a BBSRC responsive mode grant (BB/V008021/1), an MRC Proximity to Discovery EXCITEME2 grant (MCPC17189) and an award from the Gordon and Betty Moore Foundation Marine Microbiology Initiative (GBMF5514). M.V. and K.T.A. gratefully acknowledge financial support from the EPSRC via grant EP/T017856/1. K.K.L was supported via a Living Systems Institute PhD studentship. A.C. was supported via an EPSRC DTP PhD studentship (EP/M506527/1). M.R.L.S. was supported by an Australian Postgraduate Award and an Institute for Molecular Biosciences Research Advancement Award. B.Z was supported by a CSC scholarship. M.A.T.B. was supported in part by Wellcome Trust Strategic Grant WT1104797/Z/14/Z and NHMRC Development grant APP1113719. This work was further supported by a Royal Society Research Grant (RG180007) awarded to S.P., a QUEX Initiator grant awarded to S.P., K.T.A. and M.A.T.B., an NHMRC Ideas grant (2004367) awarded to M.A.T.B, and a GW4 Initiator award to M.V., K.T.A and S.P.. S.P.’s work in this area is also supported by a Marie Skłodowska-Curie project SINGEK (H2020-MSCA-ITN-2015-675752).

## Competing interests

The authors declare no competing interests.

## Author Contributions

S.P. designed the research. S.P. and U.L. developed the project. U.L., K.K.L. and A.C. performed the experiments. M.V. and K.T.A. developed and implemented the mathematical model. M.R.L.S., W.P., B.Z. and M.B. designed and synthesised the library of fluorescent antibiotic derivatives. U.L., M.V., K.K.L., A.C., M.R.L.S., W.P., B.Z. K.T.A., M.B. and S.P. analysed and discussed the data. U.L. and S.P. wrote the paper. All authors read and approved the final manuscript.

---

**Figure S1.**
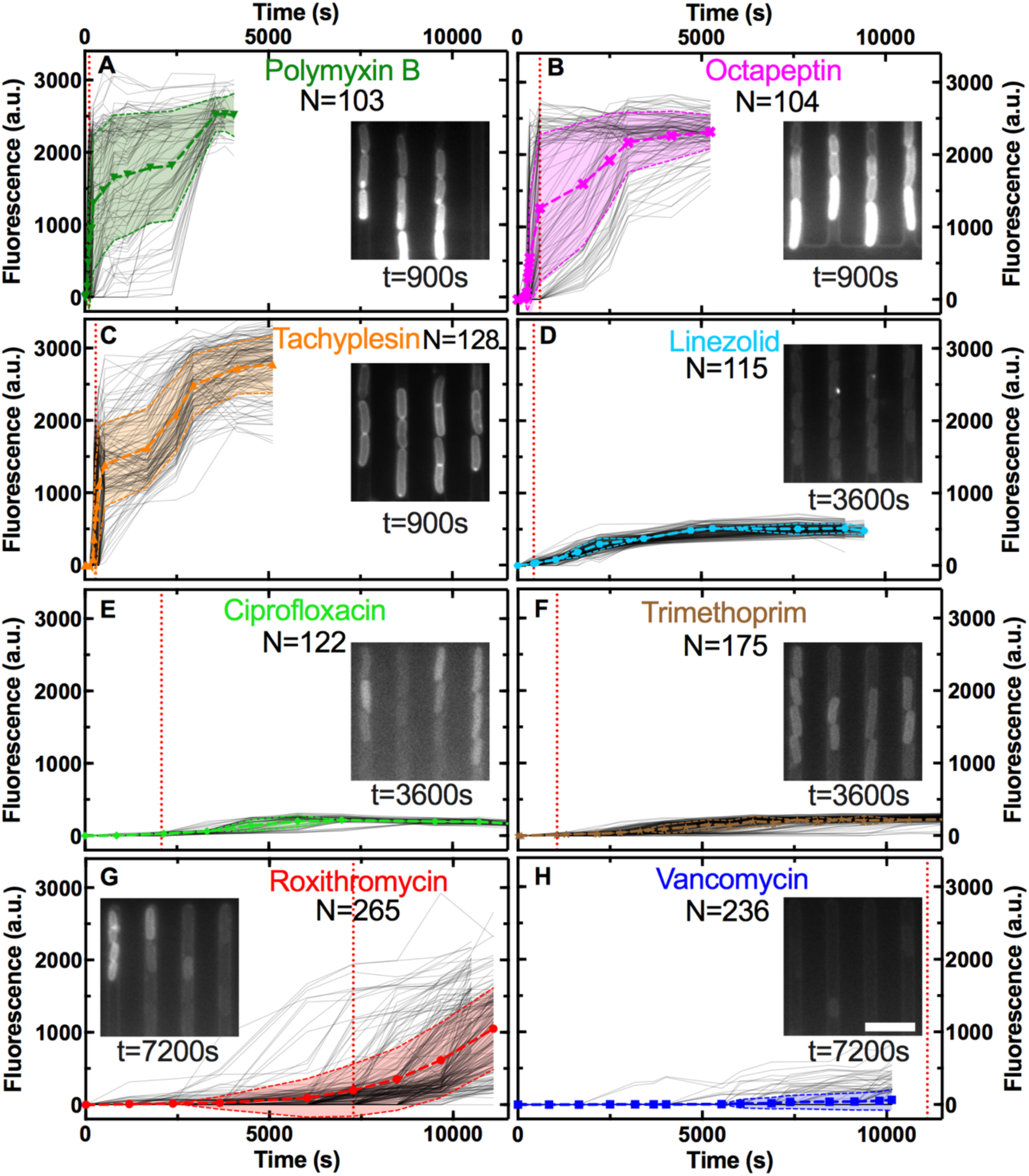
Accumulation of the fluorescent derivatives of **A**) polymyxin B, **B**) octapeptin, **C**) tachyplesin, **D**) linezolid, **E**) ciprofloxacin, **F**) trimethoprim, **G**) roxithromycin and **H**) vancomycin in 103, 104, 128, 115, 122, 175, 265 and 236 individual *E. coli*, respectively (continuous lines), after adding each probe at 46 μg mL^-1^ extracellular concentration in M9 minimal medium from t = 0 onwards. Data were collated from biological triplicate. Fluorescence values were background subtracted and normalised by cell size (see Methods). The symbols and shaded areas represent the mean and standard deviation of the corresponding single-cell values. Insets: representative fluorescence images showing the accumulation of each probe at the specific time point. Scale bar: 5 μm. The vertical dotted lines represent the time point at which the median of each dataset became larger than zero. The median remained zero throughout the entire experiments carried out with vancomycin-NBD, hence the dotted line has been arbitrarily set at 11,500 s in **H**) for comparison purposes only.

**Figure S2.**
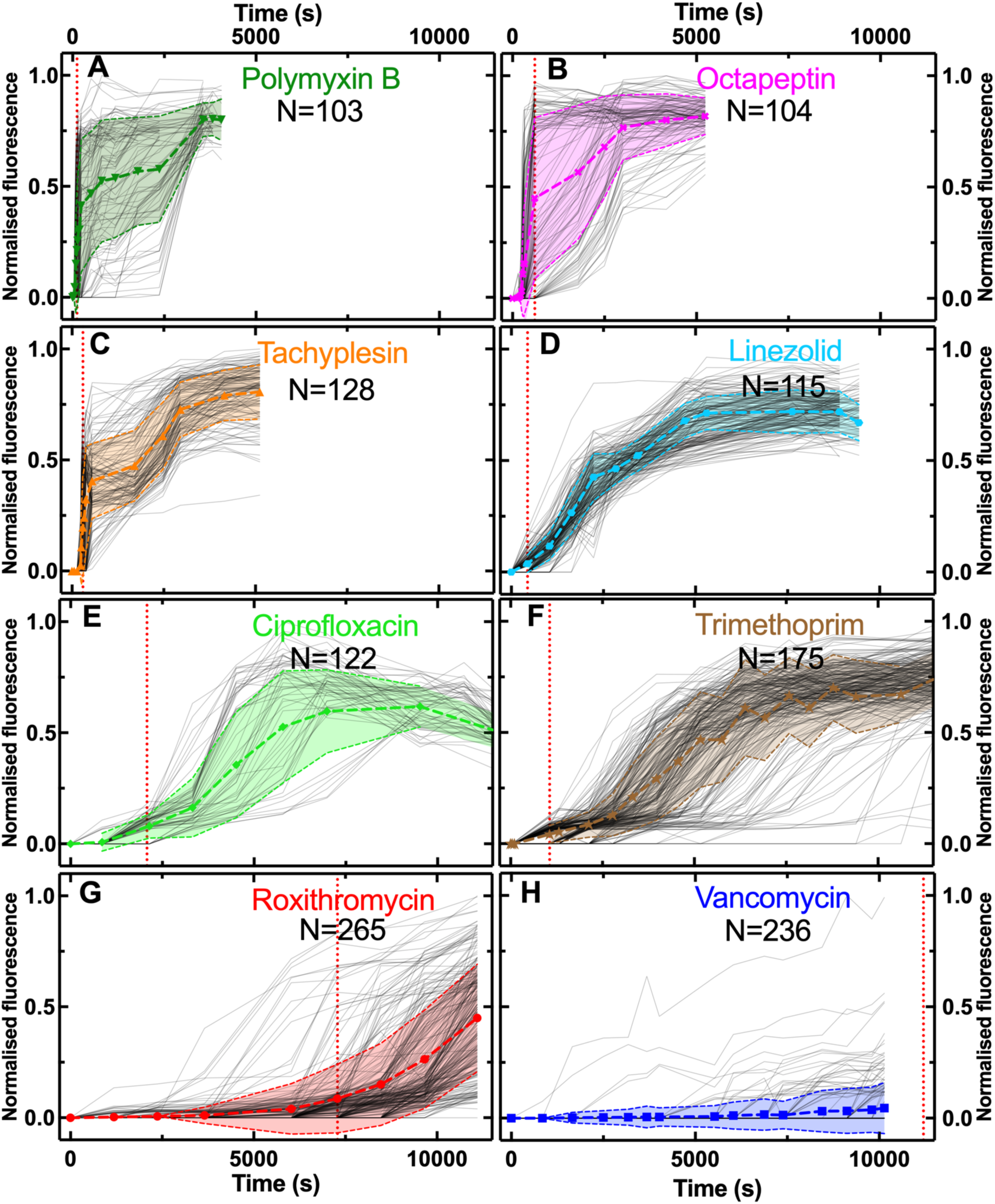
Normalised accumulation of the fluorescent derivatives of **A**) polymyxin B, **B**) octapeptin, **C**) tachyplesin, **D**) linezolid, **E**) ciprofloxacin, **F**) trimethoprim, **G**) roxithromycin and **H**) vancomycin. These data are reproduced from Fig. S1 after normalising all fluorescent values to the maximum fluorescence value in each dataset.

**Figure S3.**
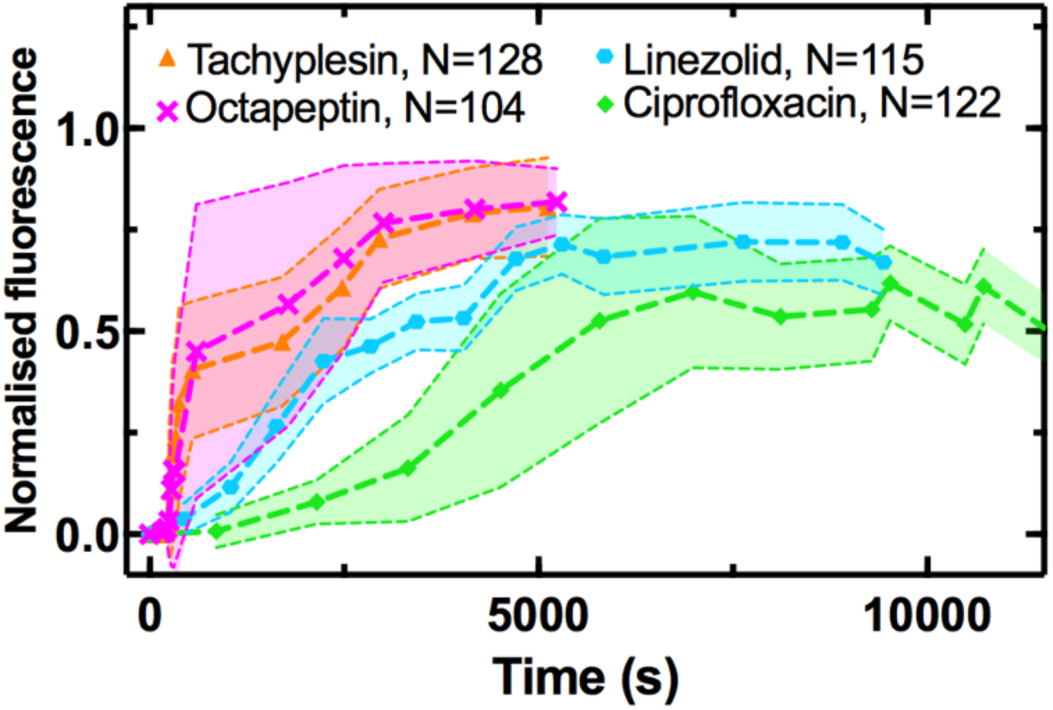
Population averages (symbols) and standard deviations (shaded areas) of the accumulation of the fluorescent derivatives of tachyplesin (triangles), octapeptin (crosses), linezolid (hexagons) and ciprofloxacin (diamonds) added at 46 μg mL^-1^ extracellular concentration in M9 minimal medium from t = 0 onwards. Data were obtained by averaging N = 128, 104, 115 and 122 single-cell values, respectively, collated from biological triplicate presented in Fig. S1.

**Figure S4.**
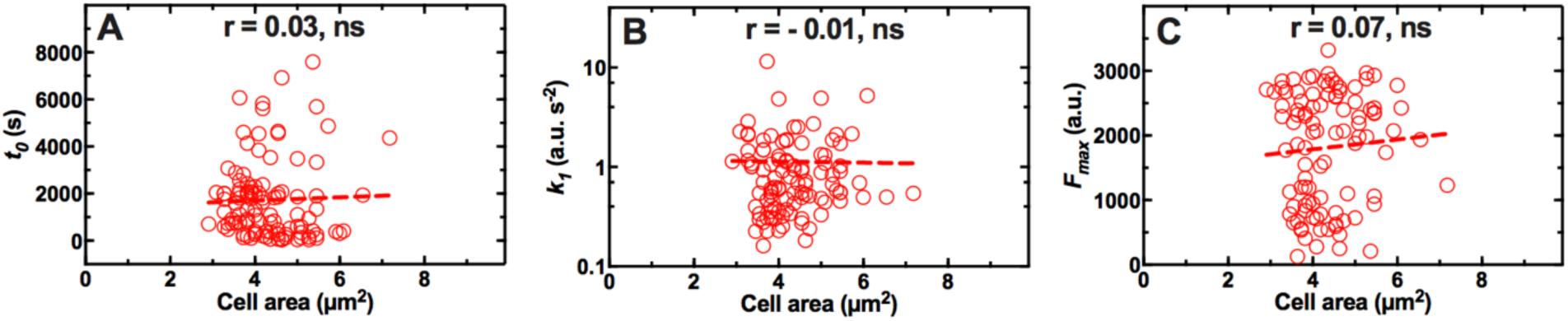
Absence of correlation between the area of each single bacterium before antibiotic treatment and the kinetic parameters **A**) *t*_*0*_, **B**) *k*_*1*_ and **C**) *F*_*max*_ describing the onset, uptake rate and level of saturation of the fluorescent derivative of roxithromycin in N = 104 *E. coli* after adding the probe at 192 μg mL^-1^ extracellular concentration in M9 minimal medium from t = 0 onwards. Data were collated from biological triplicate. We also found no correlation between cell area and the three kinetic accumulation parameters above for the other seven antibiotic probes investigated.

**Figure S5.**
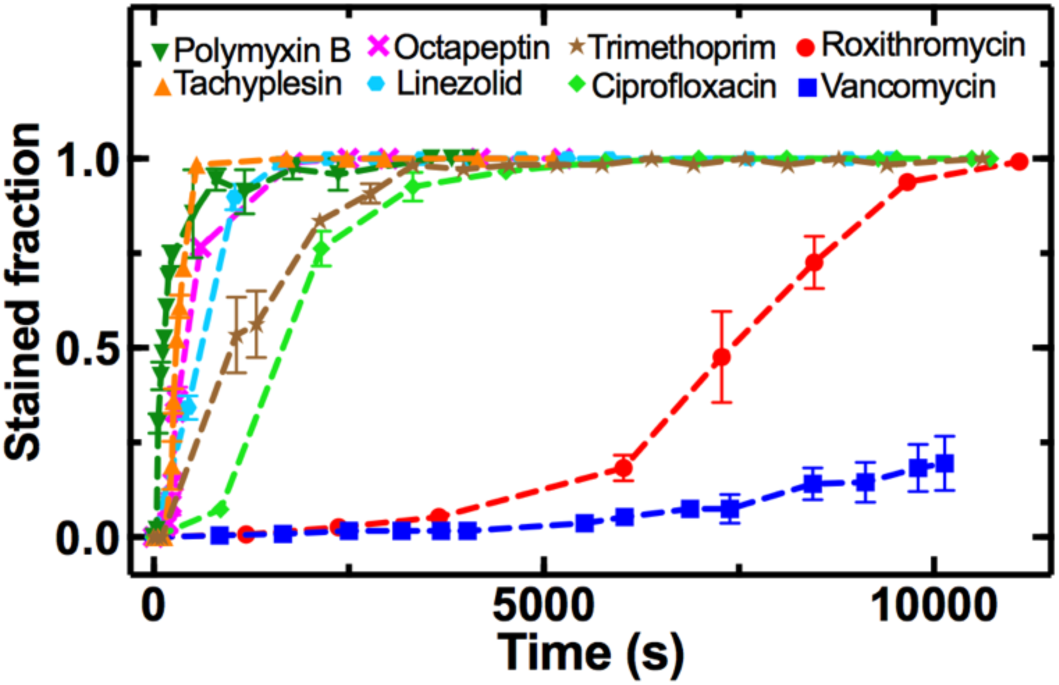
Temporal dependence of the fraction of *E. coli* stained by fluorescent derivatives of polymyxin B (downwards triangles), tachyplesin (upwards triangles), octapeptin (crosses), linezolid (hexagons), trimethoprim (stars), ciprofloxacin (diamonds), roxithromycin (circles) or vancomycin (squares). The stained fraction at each time point is defined as the ratio of the number of bacteria displaying a fluorescence distinguishable from the background over the total number of bacteria at that time point. Symbols and error bars are the mean and standard error of the mean values calculated by averaging the N = 103, 128, 104, 115, 175, 122, 265, 236 individual bacteria, respectively, from biological triplicate presented in Fig. S1.

**Figure S6.**
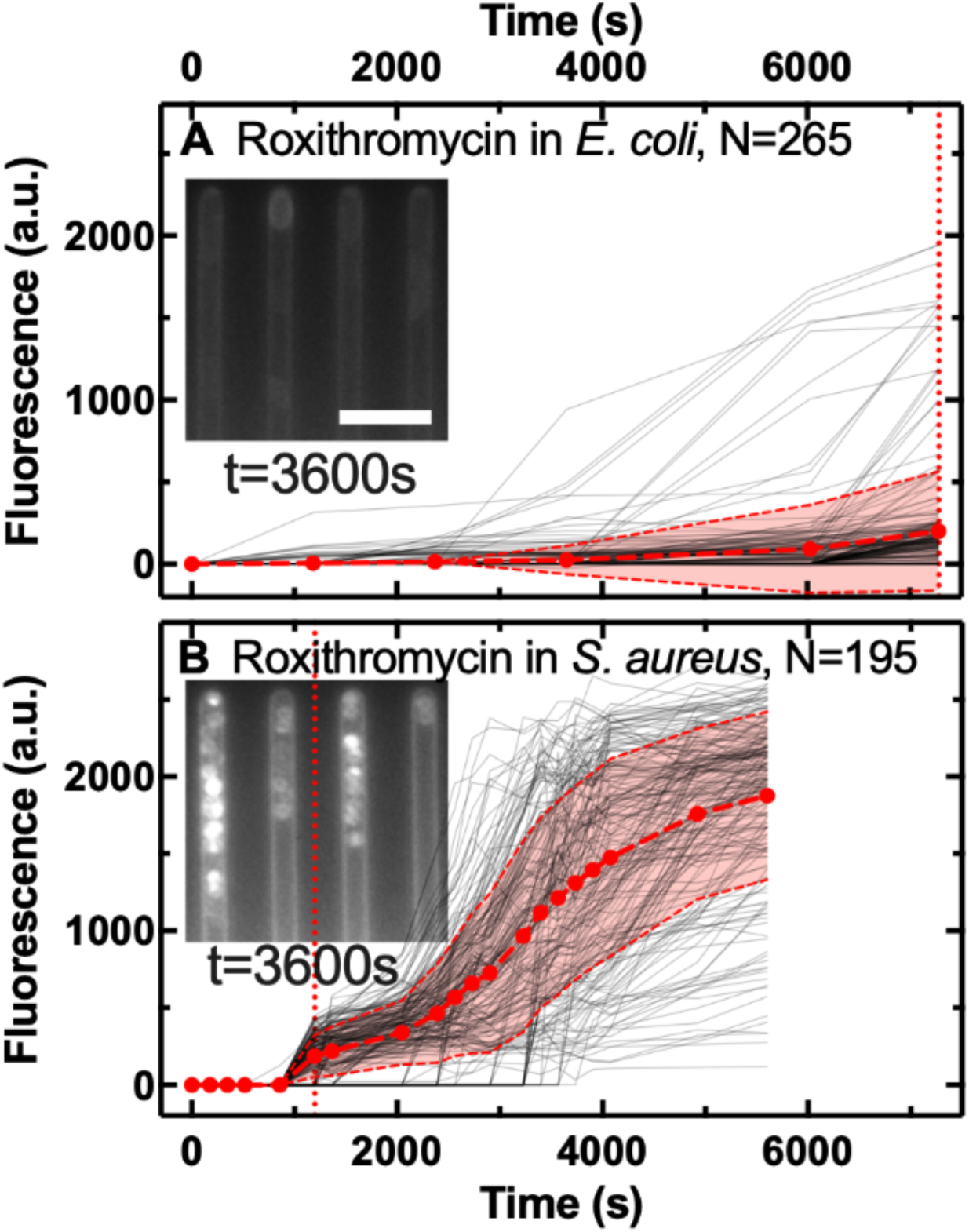
Accumulation of the fluorescent derivative of roxithromycin in **A**) N = 265 individual *E. coli* and **B**) N = 195 individual *S. aureus* (continuous lines), after adding the probe at 46 μg mL^-1^ extracellular concentration in M9 minimal medium from t = 0 onwards. Data were collated from biological triplicate. Fluorescence values were background subtracted and normalised by cell size. The symbols and shaded areas are the mean and standard deviation of the corresponding single-cell values. Insets: representative fluorescence images showing the accumulation of the fluorescent derivative of roxithromycin 3,600 s post addition to the bacteria hosting channels. Scale bar: 5 μm. The vertical dotted lines represent the time points at which the median of each dataset became larger than zero.

**Figure S7.**
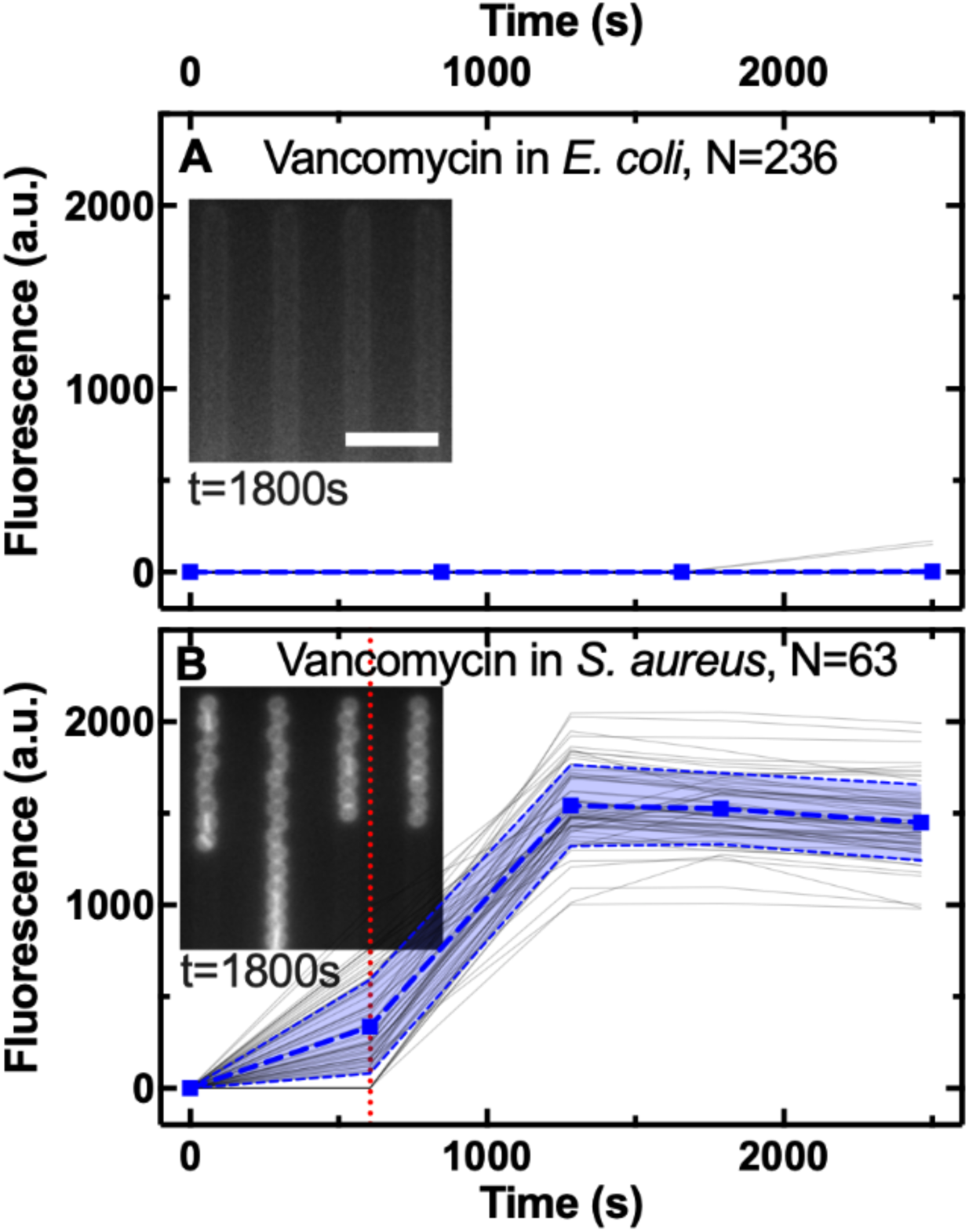
Accumulation of the fluorescent derivative of vancomycin in **A**) N = 236 individual *E. coli* and **B**) N = 63 individual *S. aureus* (continuous lines) cells, after adding the probe at 46 μg mL^-1^ extracellular concentration in M9 minimal medium from t = 0 onwards. Data were collated from biological triplicate. Fluorescence values were background subtracted and normalised by cell size. The symbols and shaded areas are the mean and standard deviation of the corresponding single-cell values. Insets: representative fluorescence images showing the accumulation of the fluorescent derivative of roxithromycin 1,800 s post addition to the bacteria hosting channels. Scale bar: 5 μm. The vertical dotted lines represent the time points at which the median of each dataset became larger than zero.

**Figure S8.**
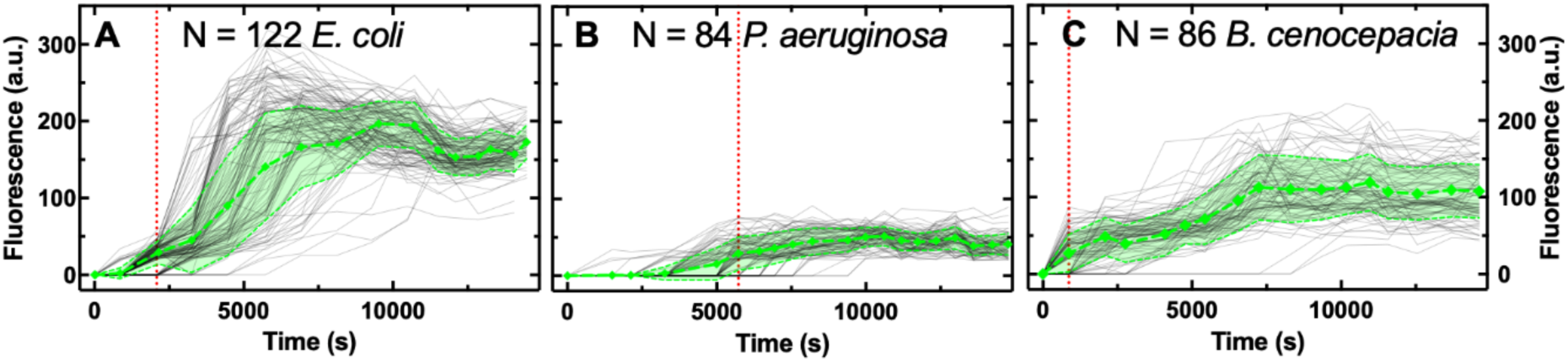
Accumulation of the fluorescent derivative of ciprofloxacin in **A**) N = 122 individual *E. coli*, **B**) N = 84 individual *P. aeruginosa* and **C**) N = 86 individual *B. cenocepacia* (continuous lines) cells, after adding the probe at 46 μg mL^-1^ extracellular concentration in M9 minimal medium from t = 0 onwards. Data were collated from biological triplicate. Fluorescence values were background subtracted and normalised by cell size. The symbols and shaded areas are the mean and standard deviation of the corresponding single-cell values. The vertical dotted lines represent the time points at which the median of each dataset became larger than zero. As expected ciprofloxacin-NBD accumulated to a significantly lower extent in *P. aeruginosa* since it lacks general porins, thus displaying a lower permeability compared to *E. coli* and *B. cenocepacia*(1).

**Figure S9.**
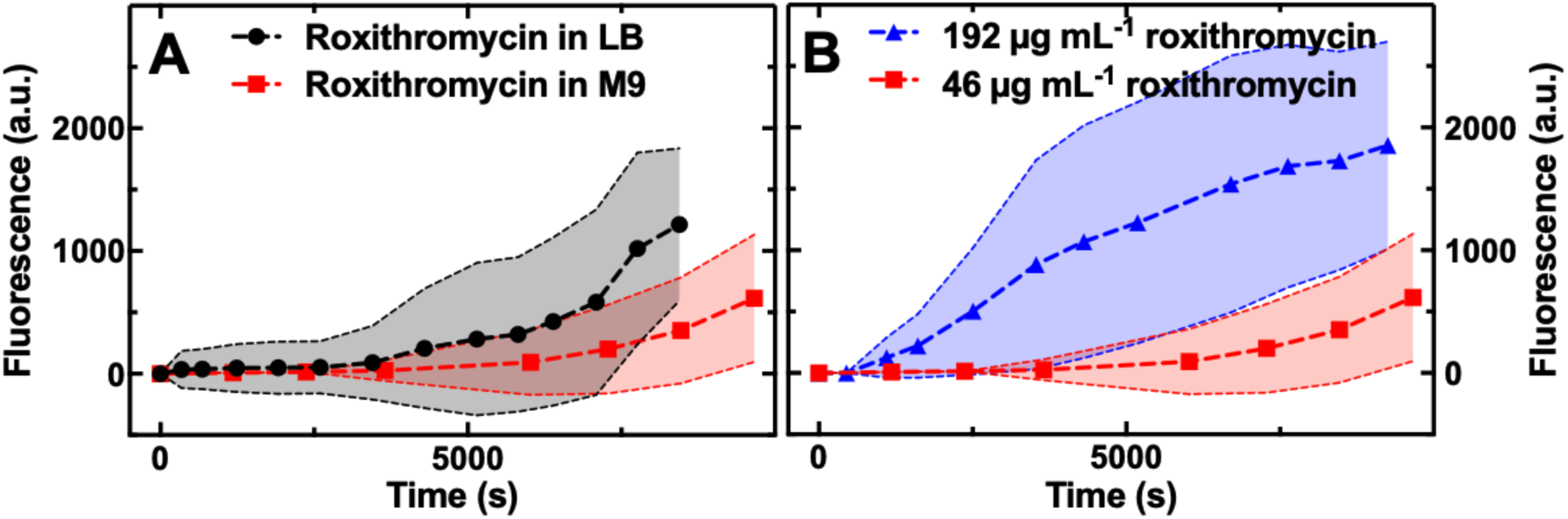
**A**) Accumulation of the fluorescent derivative of roxithromycin in LB (circles) or M9 medium (squares) drug milieu delivered to N = 46 and 265 individual *E. coli*, respectively, at an extracellular concentration of 46 μg mL^-1^. **B**) Accumulation of the fluorescent derivative of roxithromycin delivered at a concentration of 192 (triangles) and 46 (squares) μg mL^-1^ in a M9 medium drug milieu to N = 110 and 265 individual *E. coli*, respectively. In both figures data were collated from biological triplicate and fluorescence values were background subtracted and normalised by cell size. The symbols and shaded areas represent the mean and standard deviation of the corresponding single-cell values.

**Figure S10.**
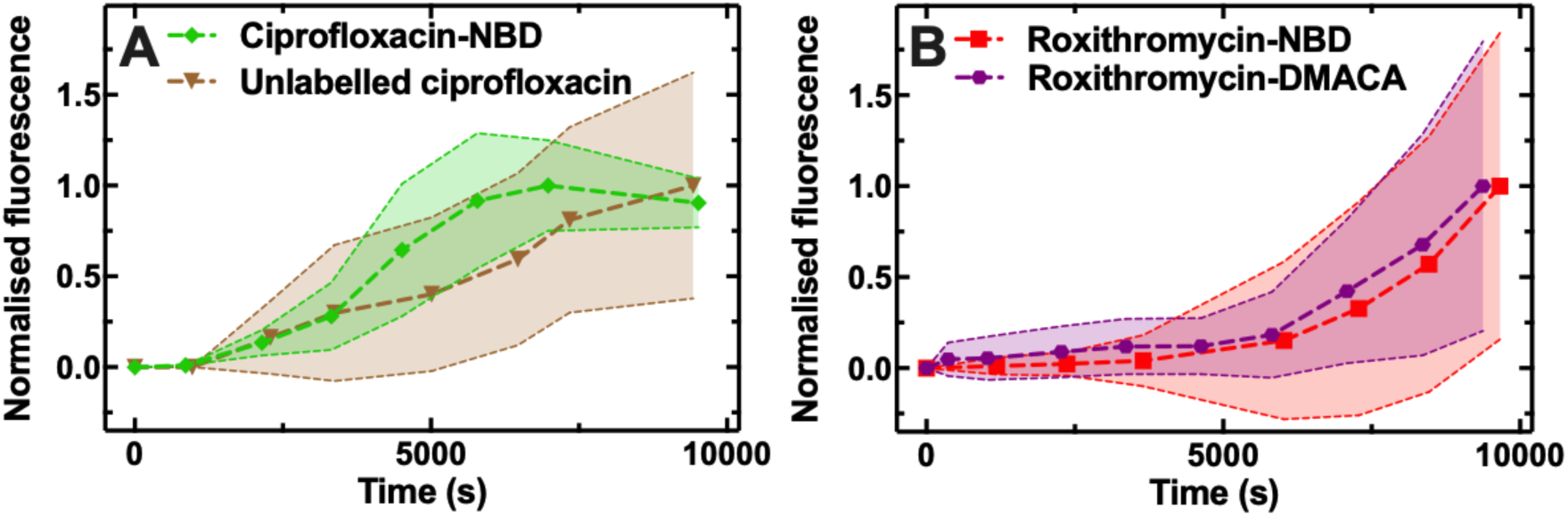
**A**) Accumulation of unlabelled ciprofloxacin (triangles) and of the fluorescent derivative ciprofloxacin-NBD (diamonds) delivered to N = 48 and 122 individual *E. coli*, respectively, at an extracellular concentration of 200 and 46 μg mL^-1^ in M9 medium, respectively. It is worth noting that unlabelled ciprofloxacin was not detectable neither extracellularly nor intracellularly at concentrations below 200 μg mL^-1^. **B**) Accumulation of the fluorescent derivatives roxithromycin-NBD (squares) and roxithromycin-DMACA (hexagons) at an extracellular concentration of 46 μg mL^-1^ in a M9 medium drug milieu delivered to N = 265 and 77 individual *E. coli*, respectively. In both figures data were collated from biological triplicate and fluorescence values were background subtracted and normalised by cell size. The symbols and shaded areas are the mean and standard deviation of the corresponding single-cell values normalised to the maximum mean fluorescence value in each dataset.

**Figure S11.**
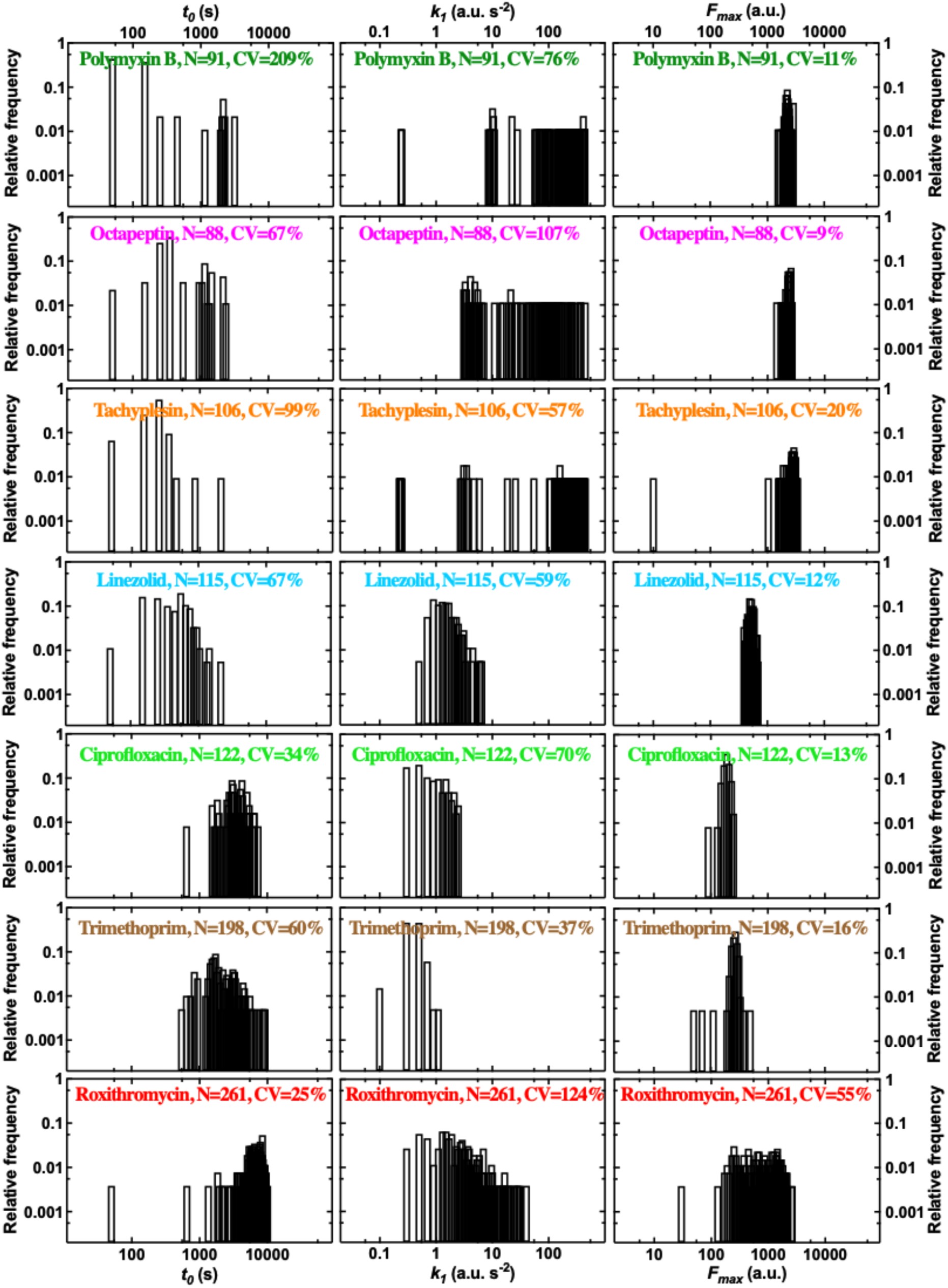
Distributions of *t*_*0*_, *k*_*1*_ and *F*_*max*_ kinetic parameters describing the accumulation of the fluorescent derivatives of polymyxin B, octapeptin, tachyplesin, linezolid, ciprofloxacin, trimethoprim and roxithromycin (from top to bottom, respectively). These parameters were inferred by fitting the single-cell data reported in Fig. S1 using our mathematical model (see Methods). Data for which the fitting algorithm returned divergent transitions were not reported and typically represented less than 1% of the data (compare N here and in Fig. S1). *t*_*0*_ is the inferred accumulation onset, i.e. the time at which each bacterium fluorescence became distinguishable from background fluorescence, *k*_*1*_ is the inferred rate of uptake, *F*_*max*_ is the inferred fluorescence saturation level at steady-state. CV is the coefficient of variation of the single-cell values in each dataset.

**Figure S12.**
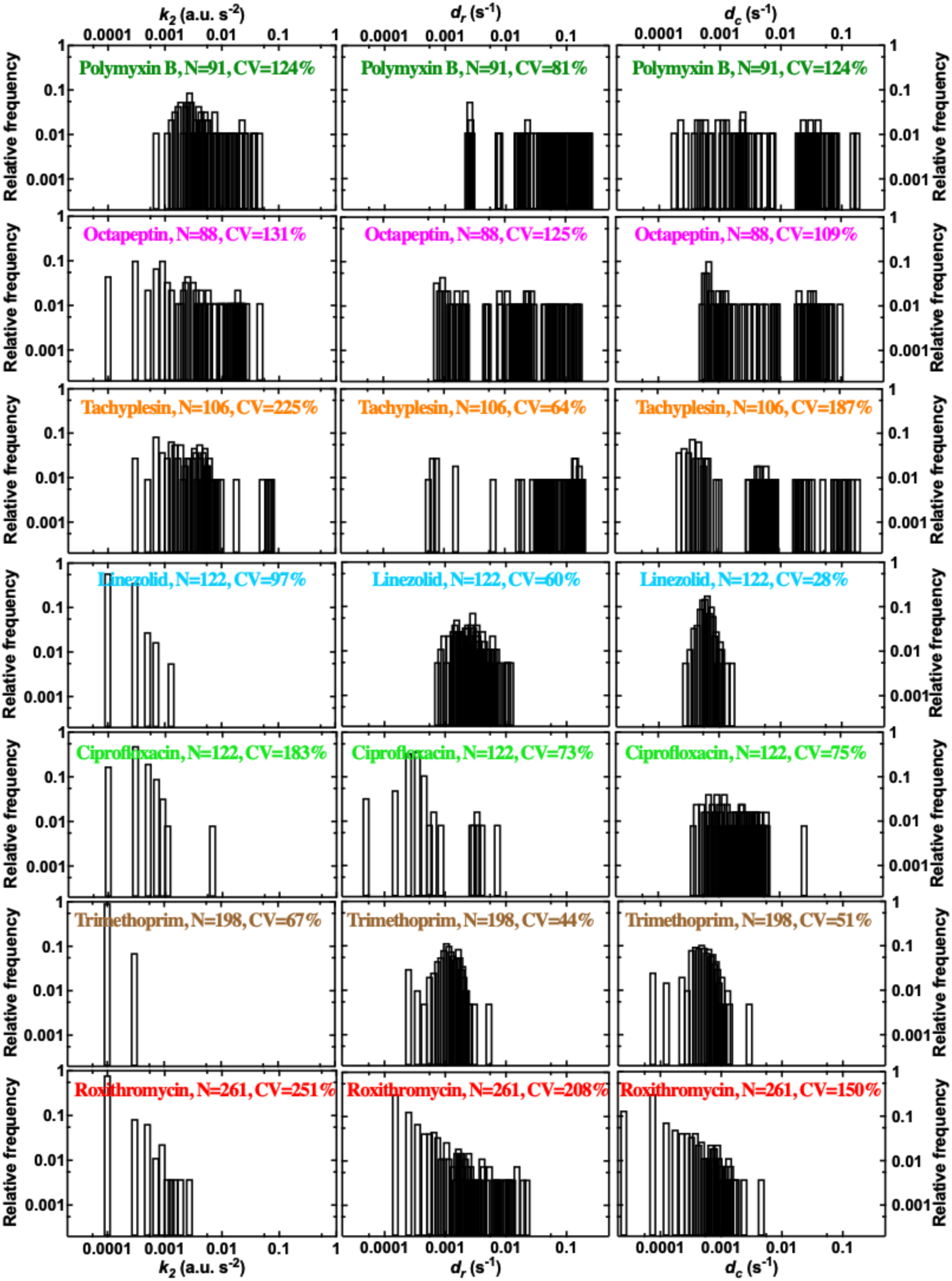
Distributions of *k*_*2*_, *d*_*r*_ and *d*_*c*_ kinetic parameters describing the accumulation of fluorescent antibiotic derivatives of polymyxin B, octapeptin, tachyplesin, linezolid, ciprofloxacin, trimethoprim and roxithromycin (from top to bottom, respectively). These parameters were inferred by fitting the single-cell data reported in Fig. S1 using our mathematical model (see Methods). Data for which the fitting algorithm returned divergent transitions were not reported and typically represented less than 1% of the data (compare N here and in Fig. S1). *k*_*2*_ is the inferred adaptive inhibitory rate constant that describes the dip we observed in some single-cell trajectories in Fig. S1, *d*_*r*_ is the drug loss rate constant, *d*_*c*_ is the dampening rate constant. CV is the coefficient of variation of the single-cell values in each dataset. Membrane targeting antibiotic probes displayed, on average, a higher adaptive inhibitory rate constant (*k*_*2*_ = 0.006, 0.007 and 0.006 a.u. s^-2^ for tachyplesin, polymyxin B and octapeptin, respectively) compared to antibiotics with intracellular targets (*k*_*2*_ = 0.0001, 0.00005, 0.0003 and 0.0001 s for linezolid, trimethoprim, ciprofloxacin and roxithromycin, respectively). Remarkably, we found notable cell-to-cell differences in *k*_*2*_ across all investigated drugs with a maximum CV of 251% for roxithromycin and a minimum CV of 67% for trimethoprim. Membrane targeting antibiotic probes also displayed, on average, a higher drug loss rate constant (*d*_*r*_ = 0.09, 0.09 and 0.03 s^-1^ for tachyplesin, polymyxin B and octapeptin, respectively) compared to antibiotics with intracellular targets (*d*_*r*_ = 0.0003, 0.001, 0.0005 and 0.001 s for linezolid, trimethoprim, ciprofloxacin and roxithromycin, respectively). Remarkably, we found notable cell-to-cell differences in *d*_*r*_ across all investigated drugs with a maximum CV of 208% for roxithromycin and a minimum CV of 44% for trimethoprim. Membrane targeting antibiotic probes also displayed, on average, a higher dampening rate constant (*d*_*c*_ = 0.009, 0.01 and 0.009 s^-1^ for tachyplesin, polymyxin B and octapeptin, respectively) compared to antibiotics with intracellular targets (*d*_*c*_ = 0.0006, 0.0005, 0.002 and 0.0003 s for linezolid, trimethoprim, ciprofloxacin and roxithromycin, respectively). Remarkably, we found notable cell-to-cell differences in *d*_*c*_ across all investigated drugs with a maximum CV of 187% for tachyplesin and a minimum CV of 28% for linezolid.

**Figure S13.**
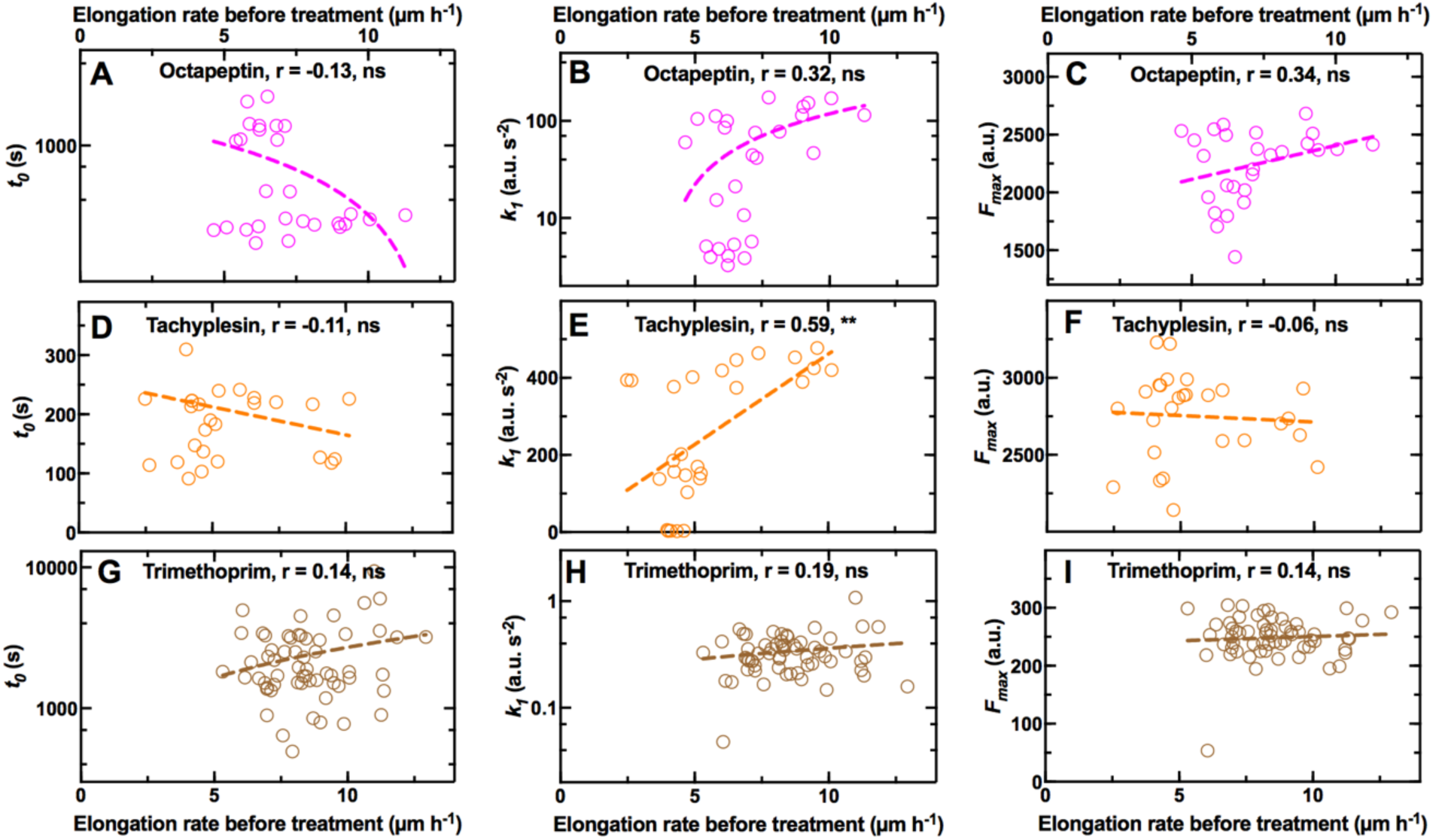
Interdependence between single-cell elongation rate before treatment and the onset *t*_*0*_, the rate *k*_*1*_, and the saturation *F*_*max*_ in the accumulation of fluorescent derivatives of **A-C**) octapeptin, **D-F**) tachyplesin and **G-I**) trimethoprim, respectively. r is the Pearson correlation coefficient, **: p-value < 0.01, ns: not significant, p-value > 0.05. N = 28, 27 and 61 individual *E. coli* investigated for the accumulation of the fluorescent derivatives of octapeptin, tachyplesin, and trimethoprim, respectively, and collated from biological triplicate. In each experiment *E. coli* were grown for 2 h in the microfluidic device with continuous supply of fresh LB. During this 2 h growth period the elongation rate of each bacterium was measured between consecutive time points and the average elongation rate for each bacterium was calculated. At the end of this 2 h growth period one of the three fluorescent antibiotic derivatives above was continuously delivered for a 4 h treatment period in the microfluidic device at a concentration of 46 μg mL^-1^ in M9 minimal medium. During this 4 h treatment period single-cell fluorescence data were obtained and dynamic accumulation parameters *t*_*0*_, *k*_*1*_ and *F*_*max*_ were inferred by fitting these single-cell data to our mathematical model (see Methods).

**Figure S14.**
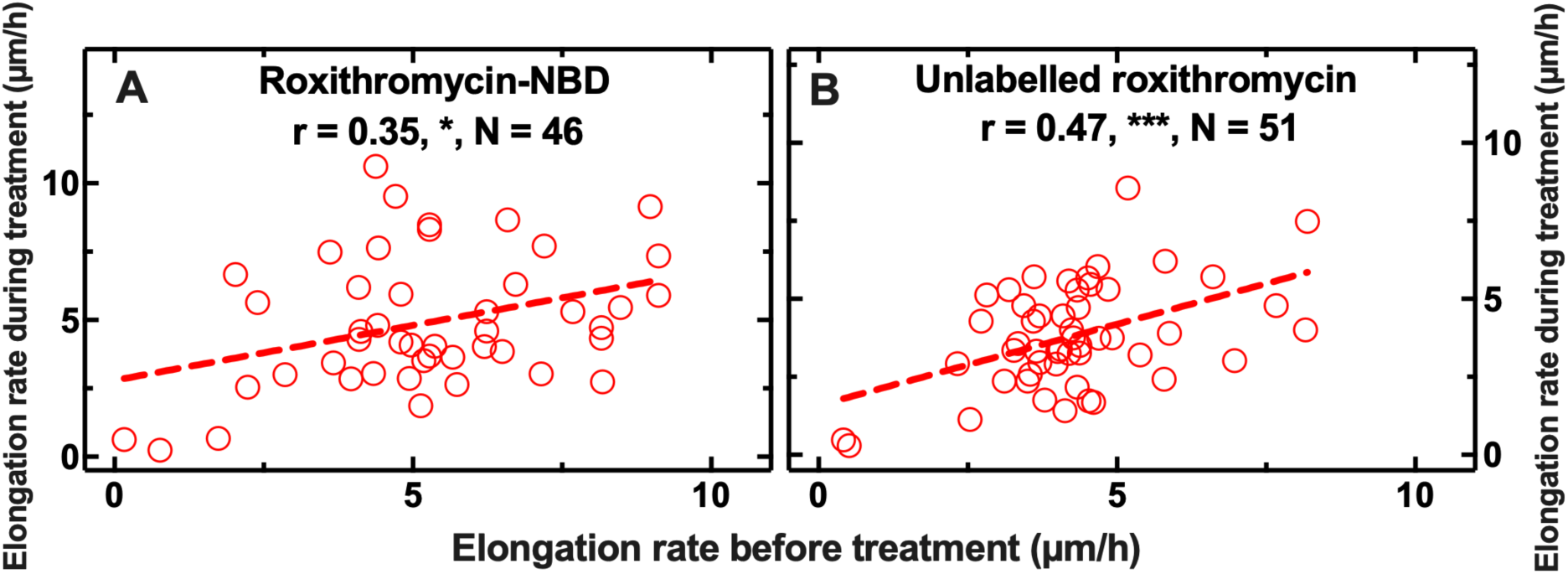
Interdependence between single-cell elongation rate before treatment and single-cell elongation rate during exposure to **A**) roxithromycin-NBD and **B**) unlabelled roxithromycin. r is the Pearson correlation coefficient, *: p-value < 0.05, ***: p-value < 0.001. N = 46 and 51 individual *E. coli* investigated and collated from biological triplicate. In each experiment *E. coli* were grown for 2 h in the microfluidic device with continuous supply of fresh LB. During this 2 h growth period the elongation rate of each bacterium was measured between consecutive time points and the average elongation rate for each bacterium was calculated. At the end of this 2 h growth period, 46 μg mL^-1^ roxithromycin-NBD or unlabelled roxithromycin dissolved in LB was continuously delivered for a 4 h treatment period in the microfluidic device. During this 4 h treatment period the elongation rate of each bacterium was measured as indicated above.

**Table S1.**
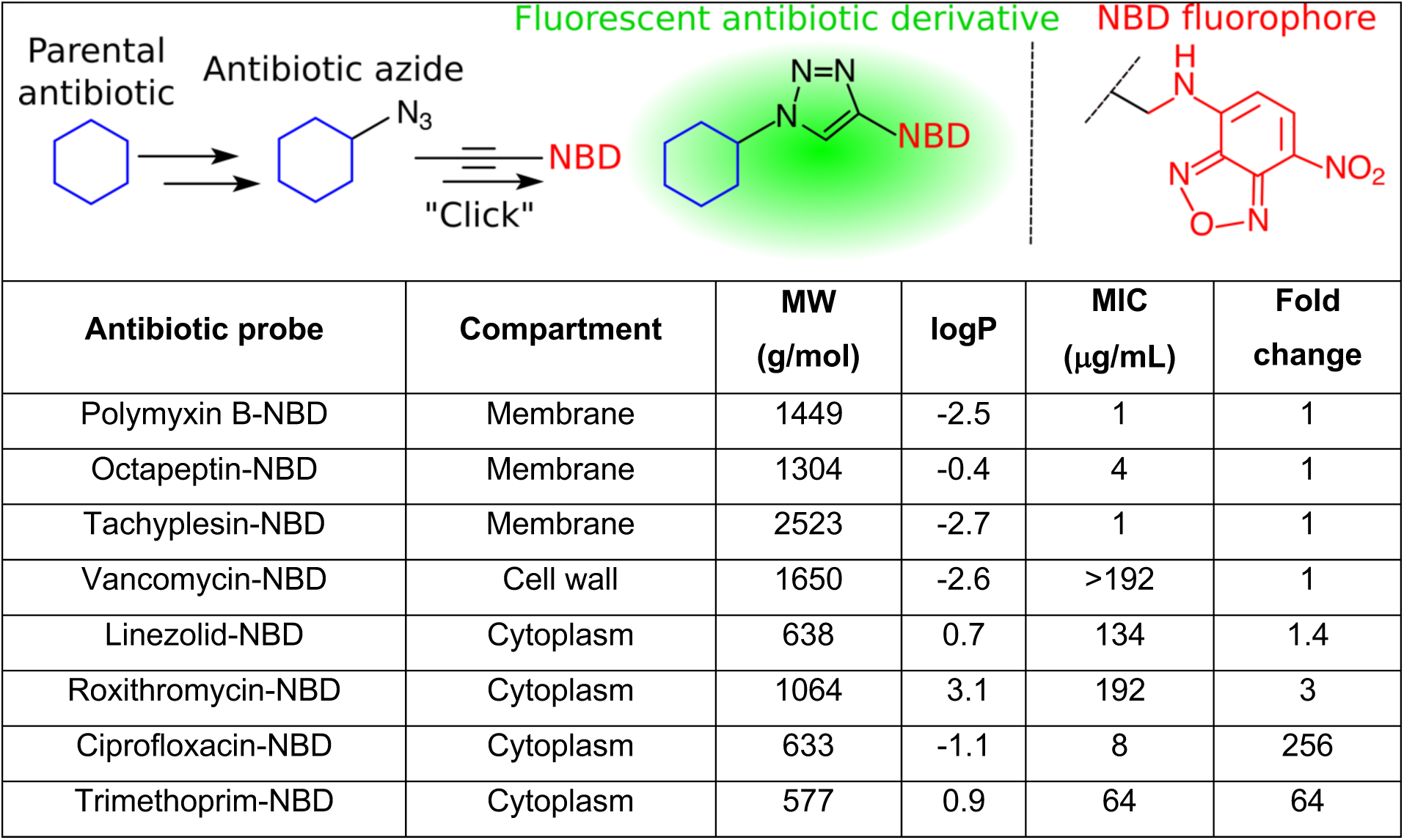
List of fluorescent antibiotic derivatives (obtained by linking the parental antibiotic to nitrobenzoxadiazole, NBD, see Methods), the bacterial compartment where their target is located, their molecular weight (MW) after linkage to NBD, their partition coefficient (logP), their measured minimum inhibitory concentration (MIC) against *E. coli* BW25113, and the fold-change compared to the MIC measured for each corresponding parental antibiotic (see Methods). MIC data were collated from biological triplicate.

**Table S2.** Pearson correlation coefficients and significance of the correlation between *t*_*0*_ and *k*_*1*_, *t*_*0*_ and *F*_*max*_ and *k*_*1*_ and *F*_*max*_ for the accumulation in single *E. coli* of all the fluorescent antibiotic derivatives investigated (apart from vancomycin) in individual *E. coli*. Data from Fig. S11 were used for these statistical comparisons. ****: p-value < 0.0001, ***: p-value < 0.001, **: p-value < 0.01, *: p-value < 0.05, ns: not significant, p-value > 0.05.

| Antibiotics | Pearson correlation coefficients and significance |  |  |
| --- | --- | --- | --- |
| | $t_0$ vs $k_1$ | $t_0$ vs $F_{max}$ | $k_1$ vs $F_{max}$ |
| Polymyxin B | -0,51, **** | -0,54, **** | 0,56, **** |
| Octapeptin | -0,46, **** | -0,61, **** | 0,20, ns |
| Tachyplesin | -0,13, ns | -0,10, ns | -0,01, ns |
| Linezolid | 0.03, ns | -0,21, ** | 0,05, ns |
| Ciprofloxacin | -0,12, ns | -0,11, ns | 0,29, *** |
| Trimethoprim | 0,06, ns | -0,32, **** | 0,11, ns |
| Roxithromycin | -0,22, *** | -0,10, ns | 0,41, **** |
| All antibiotics | -0,40, **** | -0,27, **** | 0,65, **** |

**Table S3.**
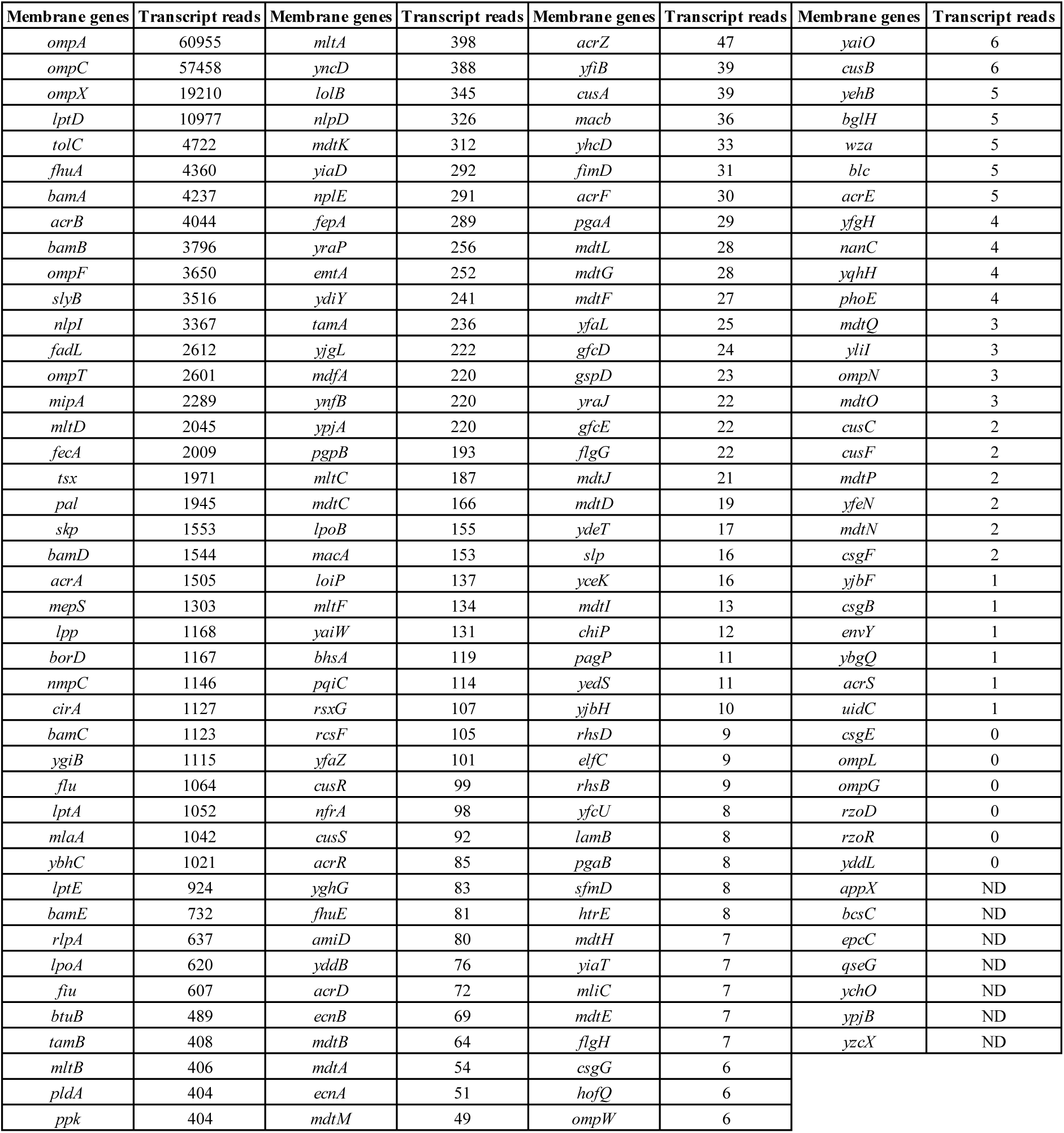
List of genes encoding outer membrane proteins (i.e. porins) and efflux pumps compiled using EcoCyc as previously reported(2), alongside their transcript reads after a 2 h growth period in LB (i.e. the time point at which antibiotic treatment starts in our microfluidic experiments) measured via RNA-sequencing as previously reported(3). Note that it has been reported that permeability of solutes through OmpA (with the most highly expressed transcripts) is a hundred fold lower compared to that through OmpC(4) (with the second most highly expressed transcripts), hence we decided to investigate the role played by OmpC in the heterogeneity in the intracellular accumulation of roxithromycin (Fig. 5E).

**Movie S1**.

Real-time accumulation of the fluorescent derivative of roxithromycin in individual *Escherichia coli* and

*Staphylococcus aureus* bacteria (top and bottom panels, respectively).

**Movie S2**.

Real-time accumulation of the fluorescent derivatives of vancomycin and roxithromycin in individual

*Staphylococcus aureus* bacteria (top and bottom panels, respectively).

## Notes

### Competing Interest Statement

The authors have declared no competing interest.

## References

1. Richards TA, Massana R, Pagliara S, Hall N. Single cell ecology. Philos Trans R Soc B Biol Sci. 2019;374:20190076.

2. Ackermann M. A functional perspective on phenotypic heterogeneity in microorganisms. Nat Rev Microbiol [Internet]. 2015;13(8):497–508. Available from: http://dx.doi.org/10.1038/nrmicro3491

3. Golding I, Paulsson J, Zawilski SM, Cox EC. Real-time kinetics of gene activity in individual bacteria. Cell. 2005;123(6):1025–36.

4. Lidstrom ME, Konopka MC. The role of physiological heterogeneity in microbial population behavior. Nat Chem Biol [Internet]. 2010;6(10):705–12. Available from: http://www.nature.com/doifinder/10.1038/nchembio.436%5Cn http://www.ncbi.nlm.nih.gov/pubmed/20852608

5. Windels EM, Michiels JE, Bergh B Van Den, Fauvart M, Michiels J. Antibiotics: Combatting Tolerance To Stop Resistance. MBio. 2019;10(5):e02095.

6. Levin-reisman I, Brauner A, Ronin I, Balaban NQ. Epistasis between antibiotic tolerance, persistence, and resistance mutations. Proc Natl Acad Sci U S A. 2019;116:14734.

7. Brauner A, Fridman O, Gefen O, Balaban NQ. Distinguishing between resistance, tolerance and persistence to antibiotic treatment. Nat Rev Microbiol [Internet]. 2016;14(5):320–30. Available from: http://www.nature.com/doifinder/10.1038/nrmicro.2016.34%5Cn http://www.ncbi.nlm.nih.gov/pubmed/27080241

8. Bamford RA, Smith A, Metz J, Glover G, Titball RW, Pagliara S. Investigating the physiology of viable but non-culturable bacteria by microfluidics and time-lapse microscopy. BMC Biol. 2017;15:121.

9. Goode O, Smith A, Łapińska U, Attrill E, Carr A, Metz J, et al. Heterologous Protein Expression Favors the Formation of Protein Aggregates in Persister and Viable but Nonculturable Bacteria. ACS Infect Dis. 2021;7:1848.

10. Goormaghtigh F, Van Melderen L. Single-cell imaging and characterization of Escherichia coli persister cells to ofloxacin in exponential cultures. Sci Adv. 2019;5(6):1–15.

11. Goode O, Smith A, Zarkan A, Cama J, Invergo BM, Belgami D, et al. Persister Escherichia coli Cells Have a Lower Intracellular pH than Susceptible Cells but Maintain Their pH in Response to. MBio. 2021;12:e00909–21.

12. Mulcahy LR, Burns JL, Lory S, Lewis K. Emergence of Pseudomonas aeruginosa strains producing high levels of persister cells in patients with cystic fibrosis. J Bacteriol. 2010;192(23):6191–9.

13. Helaine S, Cheverton AM, Watson KG, Faure LM, Matthews SA, Holden DW. Internalization of Salmonella by Macrophages Induces Formation of Nonreplicating Persisters. Science (80-). 2014;343:204–8.

14. Stapels DAC, Hill PWS, Westermann AJ, Fisher RA, Thurston TL, Saliba AE, et al. Salmonella persisters undermine host immune defenses during antibiotic treatment. Science (80-). 2018;362(6419):1156–60.

15. Baltekin Ö, Boucharin A, Tano E, Andersson DI, Elf J. Antibiotic susceptibility testing in less than 30 min using direct single-cell imaging. Proc Natl Acad Sci U S A. 2017;114(34):9170–5.

16. Shatalin K, Nuthanakanti A, Kaushik A, Shishov D, Peselis A, Shamovsky I, et al. Inhibitors of bacterial H 2 S biogenesis targeting antibiotic resistance and tolerance. Science (80-). 2021;1175(June):1169–75.

17. Rybenkov V V., Zgurskaya HI, Ganguly C, Leus I V., Zhang Z, Moniruzzaman M. The Whole Is Bigger than the Sum of Its Parts: Drug Transport in the Context of Two Membranes with Active Efflux. Chem Rev. 2021;121:5597.

18. Van Bambeke F, Barcia-Macay M, Lemaire S, Tulkens PM. Cellular pharmacodynamics and pharmacokinetics of antibiotics: Current views and perspectives. Curr Opin Drug Discov Dev. 2006;9(2):218–30.

19. Six DA, Krucker T, Leeds JA. Advances and challenges in bacterial compound accumulation assays for drug discovery. Curr Opin Chem Biol [Internet]. 2018;44:9–15. Available from: https://doi.org/10.1016/j.cbpa.2018.05.005

20. Zgurskaya HI, Rybenkov V V., Krishnamoorthy G, Leus I V. Trans-envelope multidrug efflux pumps of Gram-negative bacteria and their synergism with the outer membrane barrier. Res Microbiol [Internet]. 2018;169(7–8):351–6. Available from: https://doi.org/10.1016/j.resmic.2018.02.002

21. Pagès J-M, James CE, Winterhalter M. The porin and the permeating antibiotic: a selective diffusion barrier in Gram-negative bacteria. Nat Rev Microbiol. 2008 Dec;6(12):893–903.

22. Nestorovich EM, Danelon C, Winterhalter M, Bezrukov SM. Designed to penetrate: Time-resolved interaction of single antibiotic molecules with bacterial pores. Proc Natl Acad Sci U S A. 2002;99(15):9789–94.

23. Farmer S, Li Z, Hancock REW. Influence of outer membrane mutations on susceptibility of Escherichia coti to the dibasic macroh’de azithromycin. J Antimicrob Chemother. 1992;29:27–33.

24. Silver LL. Bioorganic & Medicinal Chemistry A Gestalt approach to Gram-negative entry. Bioorg Med Chem [Internet]. 2016;24(24):6379–89. Available from: http://dx.doi.org/10.1016/j.bmc.2016.06.044

25. Cama J, Henney AM, Winterhalter M. Breaching the Barrier: Quantifying Antibiotic Permeability across Gram-negative Bacterial Membranes. J Mol Biol [Internet]. 2019;431(18):3531–46. Available from: https://doi.org/10.1016/j.jmb.2019.03.031

26. Du D, Wang Z, James NR, Voss JE, Klimont E, Ohene-Agyei T, et al. Structure of the AcrAB-TolC multidrug efflux pump. Nature [Internet]. 2014;509(7501):512–5. Available from: http://dx.doi.org/10.1038/nature13205

27. Blair JMA, Piddock LJV. How to measure export via bacterial multidrug resistance efflux pumps. MBio. 2016;7(4):1–6.

28. Blair JMA, Webber MA, Baylay AJ, Ogbolu DO, Piddock LJV. Molecular mechanisms of antibiotic resistance. Nat Rev Microbiol [Internet]. 2015;13(1):42–51. Available from: http://dx.doi.org/10.1038/nrmicro3380

29. Fitzpatrick AWP, Llabrés S, Neuberger A, Blaza JN, Bai XC, Okada U, et al. Structure of the MacAB-TolC ABC-type tripartite multidrug efflux pump. Nat Microbiol. 2017;2(May):17070.

30. Acosta-Gutiérrez S, Ferrara L, Pathania M, Masi M, Wang J, Bodrenko I, et al. Getting Drugs into Gram-Negative Bacteria: Rational Rules for Permeation through General Porins. ACS Infect Dis. 2018;4(10):1487–98.

31. Tommasi R, Brown DG, Walkup GK, Manchester JI, Miller AA. ESKAPEing the labyrinth of antibacterial discovery. Nat Rev Drug Discov. 2015;14(8):529–42.

32. Delcour AH. Electrophysiology of bacteria. Annu Rev Microbiol. 2013;67:179–97.

33. Kojima S, Nikaido H. Permeation rates of penicillins indicate that Escherichia coli porins function principally as nonspecific channels. Proc Natl Acad Sci [Internet]. 2013;110(28):E2629–34. Available from: http://www.pnas.org/cgi/doi/10.1073/pnas.1310333110

34. Piddock LJ V, Ricci V, Asuquo AE. Quinolone accumulation by Pseudomonas aeruginosa, Staphylococcus aureus and Escherichia coli. J Antimicrob Chemother. 1999;43:61–70.

35. Asuquo AE, Piddock LJ V. Accumulation and killing kinetics of fifteen quinolones for Escherichia coli, Staphylococcus aureus and Pseudomonas aeruginosa. J Antimicrob Chemother. 1993;31:865–80.

36. Zhou Y, Joubran C, Miller-Vedam L, Isabella V, Nayar A, Tentarelli S, et al. Thinking outside the “bug”: A unique assay to measure intracellular drug penetration in Gram-negative bacteria. Anal Chem. 2015;87(7):3579–84.

37. Richter MF, Drown BS, Andrew P, Garcia A, Shirai T, Svec RL, et al. Predictive compound accumulation rules yield a broad-spectrum antibiotic. Nature [Internet]. 2017;545(7654):299–304. Available from: http://dx.doi.org/10.1038/nature22308

38. Davis TD, Gerry CJ, Tan DS. General platform for systematic quantitative evaluation of small-molecule permeability in bacteria. ACS Chem Biol. 2014;9(11):2535–44.

39. Prochnow H, Fetz V, Hotop S-K, Rivera MG, Heumann A, Brönstrup M. Subcellular quantification of uptake in Gram-negative bacteria. Anal Chem [Internet]. 2019;91:1863. Available from: http://pubs.acs.org

40. Brochado AR, Telzerow A, Bobonis J, Banzhaf M, Mateus A, Selkrig J, et al. Species-specific activity of antibacterial drug combinations. Nature. 2018;559(7713):259–63.

41. Iyer R, Ye Z, Ferrari A, Duncan L, Tanudra MA, Tsao H, et al. Evaluating LC-MS/MS to measure accumulation of compounds within bacteria. ACS Infect Dis. 2018;4:1336–45.

42. Tian H, Six DA, Krucker T, Leeds JA, Winograd N. Subcellular Chemical Imaging of Antibiotics in Single Bacteria Using C60-Secondary Ion Mass Spectrometry. Anal Chem. 2017;89(9):5050–7.

43. Heidari-Torkabadi H, Che T, Lombardo MN, Wright DL, Anderson AC, Carey PR. Measuring propargyl-linked drug populations inside bacterial cells, and their interaction with a dihydrofolate reductase target, by Raman microscopy. Biochemistry. 2015;54(17):2719–26.

44. Vergalli J, Dumont E, Pajović J, Cinquin B, Maigre L, Masi M, et al. Spectrofluorimetric quantification of antibiotic drug concentration in bacterial cells for the characterization of translocation across bacterial membranes. Nat Protoc. 2018;13(6):1348–61.

45. Vergalli J, Dumont E, Cinquin B, Maigre L, Pajovic J, Bacqué E, et al. Fluoroquinolone structure and translocation flux across bacterial membrane. Sci Rep. 2017;7(1):9821.

46. Vergalli J, Atzori A, Pajovic J, Dumont E, Malloci G, Masi M, et al. The challenge of intracellular antibiotic accumulation, a function of fluoroquinolone influx versus bacterial efflux. Commun Biol. 2020;3(1):1–12.

47. Lapinska U, Glover G, Capilla-lasheras P, Young AJ, Pagliara S. Bacterial ageing in the absence of external stressors. Philos Trans R Soc B Biol Sci. 2019;374:20180442.

48. Cama J, Voliotis M, Metz J, Smith A, Iannucci J, Keyser UF, et al. Single-cell microfluidics facilitates the rapid quantification of antibiotic accumulation in Gram-negative bacteria. Lab Chip [Internet]. 2020;20(15):2765–75. Available from: http://dx.doi.org/10.1039/D0LC00242A

49. Stone MRL, Butler MS, Phetsang W, Cooper MA, Blaskovich MAT. Fluorescent Antibiotics: New Research Tools to Fight Antibiotic Resistance. Trends Biotechnol [Internet]. 2018;36(5):523–36. Available from: http://dx.doi.org/10.1016/j.tibtech.2018.01.004

50. Blaskovich MA, Phetsang W, Stone MRL, Lapinska U, Pagliara S, Bhalla R, et al. Antibiotic-derived molecular probes for bacterial imaging. In: Proceedings of SPIE - The International Society for Optical Engineering. 2019.

51. Lin L, Du Y, Song J, Wang W, Yang C. Imaging Commensal Microbiota and Pathogenic Bacteria in the Gut. Acc Chem Res. 2021;54(9):2076–87.

52. Stone MRL, Łapińska U, Pagliara S, Masi M, Blanchfield JT, Cooper MA, et al. Fluorescent macrolide probes – synthesis and use in evaluation of bacterial resistance. RSC Chem Biol. 2020;1:395–404.

53. Phetsang W, Blaskovich MAT, Butler MS, Huang JX, Zuegg J, Mamidyala SK, et al. An azido-oxazolidinone antibiotic for live bacterial cell imaging and generation of antibiotic variants. Bioorganic Med Chem [Internet]. 2014;22(16):4490–8. Available from: http://dx.doi.org/10.1016/j.bmc.2014.05.054

54. Blaskovich MA, Phetsang W, Stone MR, Lapinska U, Pagliara S, Bhalla R, et al. Antibiotic-derived molecular probes for bacterial imaging. In: Photonic Diagnosis and Treatment of Infections and Inflammatory Diseases II [Internet]. 2019. p. 2. Available from: https://www.spiedigitallibrary.org/conference-proceedings-of-spie/10863/2507329/Antibiotic-derived-molecular-probes-for-bacterial-imaging/10.1117/12.2507329.full

55. Stone MRL, Masi M, Phetsang W, Pages J-M, Cooper MA, Blaskovich MAT. Fluoroquinolone-derived fluorescent probes for studies of bacterial penetration and efflux. Medchemcomm. 2019;10:901.

56. Phetsang W, Pelingon R, Butler MS, Kc S, Pitt ME, Kaeslin G, et al. Fluorescent Trimethoprim Conjugate Probes to Assess Drug Accumulation in Wild Type and Mutant Escherichia coli. ACS Infect Dis. 2016;2(10):688–701.

57. Silander OK, Nikolic N, Zaslaver A, Bren A, Kikoin I, Alon U, et al. A genome-wide analysis of promoter-mediated phenotypic noise in Escherichia coli. PLoS Genet. 2012;8(1).

58. Windels EM, Michiels JE, Fauvart M, Wenseleers T, Bergh B Van Den, Michiels J. Bacterial persistence promotes the evolution of antibiotic resistance by increasing survival and mutation rates. ISME J [Internet]. 2019; Available from: http://dx.doi.org/10.1038/s41396-019-0344-9

59. Pu Y, Zhao Z, Li Y, Zou J, Ma Q, Zhao Y, et al. Enhanced Efflux Activity Facilitates Drug Tolerance in Dormant Bacterial Cells. Mol Cell. 2016;62(2):284–94.

60. Taniguchi Y, Choi PJ, Li GW, Chen H, Babu M, Hearn J, et al. Quantifying E. coli proteome and transcriptome with single-molecule sensitivity in single cells. Science (80-) [Internet]. 2010;329:533–8. Available from: http://www.sciencemag.org/cgi/doi/10.1126/science.1188308

61. Murray PR. Manual of clinical microbiology. American Society for Microbiology; 1995.

62. Ude J, Tripathi V, Buyck JM, Söderholm S, Cunrath O, Fanous J, et al. Outer membrane permeability: Antimicrobials and diverse nutrients bypass porins in Pseudomonas aeruginosa. Proc Natl Acad Sci U S A. 2021;118(31):1–8.

63. Peterson AA, Fesik SW, McGroarty EJ. Decreased binding of antibiotics to lipopolysaccharide from polymyxin-resistant strains of Escherichia coli and Salmonella typhimurium. Antimicrob Agents Chemother. 1987;31(2):230–7.

64. Walters III MC, Roe F, Bugnicourt A, Franklin MJ, Stewart PS. Contributions of Antibiotic Penetration, Oxygen Limitation. Antimicrob Agents Chemother. 2003;47(1):317–23.

65. Wang P, Robert L, Pelletier J, Dang WL, Taddei F, Wright A, et al. Robust growth of Escherichia coli. Curr Biol [Internet]. 2010;20(12):1099–103. Available from: http://dx.doi.org/10.1016/j.cub.2010.04.045

66. Balaban NQ, Merrin J, Chait R, Kowalik L, Leibler S. Bacterial Persistence as a Phenotypic Switch. Science (80-) [Internet]. 2004;305(September):1622–5. Available from: http://www.ncbi.nlm.nih.gov/pubmed/15308767%5Cnhttp://www.sciencemag.org/content/305/5690/1622.short

67. Balaban NQ, Gerdes K, Lewis K, McKinney JD. A problem of persistence: still more questions than answers? Nat Rev Microbiol [Internet]. 2013;11(8):587–91. Available from: http://www.ncbi.nlm.nih.gov/pubmed/24020075

68. Lewis K. Persister cells, dormancy and infectious disease. Nat Rev Microbiol [Internet]. 2007;5(1):48–56. Available from: http://www.ncbi.nlm.nih.gov/pubmed/17143318

69. Smith A, Kaczmar A, Bamford RA, Smith C, Frustaci S, Kovacs-Simon A, et al. The culture environment influences both gene regulation and phenotypic heterogeneity in Escherichia coli. Front Microbiol. 2018;9:1739.

70. Nikaido H. Molecular basis of bacterial outer membrane permeability revisited. Microbiol Mol Biol Rev [Internet]. 2003;67(4):593–656. Available from: http://www.pubmedcentral.nih.gov/articlerender.fcgi?artid=309051&tool=pmcentrez&rendertype=abstract

71. Otto G. An arresting antitoxin. Nat Rev Microbiol [Internet]. 2021;2:41579. Available from: http://dx.doi.org/10.1038/s41579-021-00512-z

72. Vaara M. Agents That Increase the Permeability of the Outer. Microbiol Rev. 1992;56(3):395–411.

73. Balaban NQ, Helaine S, Camilli A, Collins JJ, Ghigo J-M, Hardt W-D, et al. Definitions and guidelines for research on antibiotic persistence. Nat Rev Microbiol [Internet]. 2019;17:441. Available from: http://dx.doi.org/10.1038/s41579-019-0196-3

74. Pontes MH, Groisman EA. A physiological basis for nonheritable antibiotic resistance. MBio. 2020;11(3):1–13.

75. Orman MA, Brynildsen MP. Dormancy is not necessary or sufficient for bacterial persistence. Antimicrob Agents Chemother. 2013;57(7):3230–9.

76. Peyrusson F, Varet H, Nguyen TK, Legendre R, Sismeiro O, Coppée JY, et al. Intracellular Staphylococcus aureus persisters upon antibiotic exposure. Nat Commun [Internet]. 2020;11(1):2200. Available from: http://dx.doi.org/10.1038/s41467-020-15966-7

77. Scott M, Gunderson CW, Mateescu EM, Zhang Z, Hwa T. Interdependence of Cell Growth Origins and Consequences. Science (80-). 2010;330(November):1099–102.

78. Dinos GP, Connell SR, Nierhaus KH, Kalpaxis DL. Erythromycin, roxithromycin, and clarithromycin: Use of slow-binding kinetics to compare their in vitro interaction with a bacterial ribosomal complex active in peptide bond formation. Mol Pharmacol. 2003;63(3):617–23.

79. Greulich P, Scott M, Evans MR, Allen RJ. Growth-dependent bacterial susceptibility to ribosome-targeting antibiotics. Mol Syst Biol. 2015;11:796.

80. Dai X, Zhu M, Warren M, Balakrishnan R, Patsalo V, Okano H, et al. Reduction of translating ribosomes enables Escherichia coli to maintain elongation rates during slow growth. Nat Microbiol. 2016;2(December 2016):16231.

81. Wilmaerts D, Windels EM, Verstraeten N, Michiels J. General Mechanisms Leading to Persister Formation and Awakening. Trends Genet [Internet]. 2019;1–11. Available from: https://doi.org/10.1016/j.tig.2019.03.007

82. Gollan B, Grabe G, Michaux C, Helaine S. Bacterial Persisters and Infection: Past, Present, and Progressing. Annu Rev ofMicrobiology. 2019;73:359.

83. Defraine V, Fauvart M, Michiels J. Fighting bacterial persistence: Current and emerging antipersister strategies and therapeutics Valerie. Drug Resist Updat [Internet]. 2018;38:12. Available from: https://doi.org/10.1016/j.drup.2018.03.002

84. Delcour AH. Outer Membrane Permeability and Antibiotic Resistance. Biochim Biophys Acta. 2009;1794(5):808–16.

85. Clark D. Novel antibiotic hypersensitive mutants of Escherichia coli genetic mapping and chemical characterization. FEMS Microbiol Lett. 1984;21(2):189–95.

86. Vaara M. Outer membrane permeability barrier to azithromycin, clarithromycin, and roxithromycin in gram-negative enteric bacteria. Antimicrob Agents Chemother. 1993;37(2):354–6.

87. Wu H, Moser C, Wang HZ, Høiby N, Song ZJ. Strategies for combating bacterial biofilm infections. Int J Oral Sci. 2015;7(July 2014):1–7.

88. Kepiro IE, Marzuoli I, Hammond K, Ba X, Lewis H, Shaw M, et al. Engineering Chirally Blind Protein Pseudocapsids into Antibacterial Persisters. ACS Nano. 2020;14:1609.

89. Hammond K, Cipcigan F, Al Nahas K, Losasso V, Lewis H, Cama J, et al. Switching Cytolytic Nanopores into Antimicrobial Fractal Ruptures by a Single Side Chain Mutation. ACS Nano. 2021;

90. Nonejuie P, Burkart M, Pogliano K, Pogliano J. Bacterial cytological profiling rapidly identifies the cellular pathways targeted by antibacterial molecules. Proc Natl Acad Sci U S A. 2013;110(40):16169–74.

91. Stokes JM, Yang K, Swanson K, Jin W, Cubillos-Ruiz A, Donghia NM, et al. A Deep Learning Approach to Antibiotic Discovery. Cell [Internet]. 2020;180(4):688-702.e13. Available from: https://doi.org/10.1016/j.cell.2020.01.021

92. Zaslaver A, Bren A, Ronen M, Itzkovitz S, Kikoin I, Shavit S, et al. A comprehensive library of fluorescent transcriptional reporters for Escherichia coli. Nat Methods. 2006;3(8):623–8.

93. Henry TC, Brynildsen MP. Development of Persister-FACSeq: a method to massively parallelize quantification of persister physiology and its heterogeneity. Sci Rep [Internet]. 2016;6(April):25100. Available from: http://www.nature.com/articles/srep25100

94. Blaskovich MAT, Hansford KA, Gong Y, Butler MS, Muldoon C, Huang JX, et al. Protein-inspired antibiotics active against vancomycin- and daptomycin-resistant bacteria. Nat Commun [Internet]. 2018;9(1):22. Available from: http://dx.doi.org/10.1038/s41467-017-02123-w

95. Gallardo-Godoy A, Muldoon C, Becker B, Elliott AG, Lash LH, Huang JX, et al. Activity and Predicted Nephrotoxicity of Synthetic Antibiotics Based on Polymyxin B. J Med Chem. 2016;59(3):1068–77.

96. Velkov T, Gallardo-Godoy A, Swarbrick JD, Blaskovich MAT, Elliott AG, Han M, et al. Structure, Function, and Biosynthetic Origin of Octapeptin Antibiotics Active against Extensively Drug-Resistant Gram-Negative Bacteria. Cell Chem Biol. 2018;25(4):380-391.e5.

97. Edwards IA, Elliott AG, Kavanagh AM, Blaskovich MAT, Cooper MA. Structure-Activity and â’Toxicity Relationships of the Antimicrobial Peptide Tachyplesin-1. ACS Infect Dis. 2017;3(12):917–26.

98. Locatelli E, Pierno M, Baldovin F, Orlandini E, Tan Y, Pagliara S. Single-File Escape of Colloidal Particles from Microfluidic Channels. Phys Rev Lett [Internet]. 2016;117(3):038001. Available from: http://link.aps.org/doi/10.1103/PhysRevLett.117.038001

99. Cama J, Pagliara S. Microfluidic Single-Cell Phenotyping of the Activity of Peptide-Based Antimicrobials. In: Polypeptide Materials: Methods and Protocols Methods in Molecular Biology. 2021. p. 237–53.

100. Smith A, Metz J, Pagliara S. MMHelper: An automated framework for the analysis of microscopy images acquired with the mother machine. Sci Rep. 2019;9(1).

101. Carpenter B, Gelman A, Hoffman MD, Lee D, Goodrich B, Betancourt M, et al. Stan: A probabilistic programming language. J Stat Softw. 2017;76(1).

## References

1. V. V. Rybenkov, et al., The Whole Is Bigger than the Sum of Its Parts: Drug Transport in the Context of Two Membranes with Active Efflux. Chem. Rev. 121, 5597 (2021).

2. K. E. Kortright, B. K. Chan, P. E. Turner, High-throughput discovery of phage receptors using transposon insertion sequencing of bacteria. Proc. Natl. Acad. Sci. U. S. A. 117, 18670–18679 (2020).

3. A. Smith, et al., The culture environment influences both gene regulation and phenotypic heterogeneity in Escherichia coli. Front. Microbiol. 9, 1739 (2018).

4. E. Sugawara, H. Nikaido, Pore-forming activity of OmpA protein of Escherichia coli. J. Biol. Chem. 267, 2507–2511 (1992).

